# Endothelial Cell SMAD6 Balances ACVRL1/Alk1 Function to Regulate Adherens Junctions and Hepatic Vascular Development

**DOI:** 10.1101/2023.03.23.534007

**Authors:** Molly R Kulikauskas, Morgan Oatley, Tianji Yu, Ziqing Liu, Lauren Matsumura, Elise Kidder, Dana Ruter, Victoria L Bautch

**Affiliations:** Cell Biology and Physiology Curriculum; Department of Biology; McAllister Heart Institute; Lineberger Comprehensive Cancer Center, The University of North Carolina, Chapel Hill, NC USA

**Author notes:** Corresponding author: Victoria L. Bautch, PhD Department of Biology, CB No. 3280 The University of North Carolina at Chapel Hill Chapel Hill, NC 27599 USA. current address: Department of Physiology, Medical College of Wisconsin, Wauwatosa, WI 53226 USA.

**Keywords:** SMAD6, ALK1/ACVRL1, BMP, endothelial cells, liver sinusoid, PI3K, adherens junctions, contractility, liver development, vascular barrier

## Abstract

BMP signaling is critical to blood vessel formation and function, but how pathway components regulate vascular development is not well-understood. Here we find that inhibitory SMAD6 functions in endothelial cells to negatively regulate ALK1/ACVRL1-mediated responses, and it is required to prevent vessel dysmorphogenesis and hemorrhage in the embryonic liver vasculature. Reduced *Alk1* gene dosage rescued embryonic hepatic hemorrhage and microvascular capillarization induced by *Smad6* deletion in endothelial cells *in vivo*. At the cellular level, co-depletion of Smad6 and Alk1 rescued the destabilized junctions and impaired barrier function of endothelial cells depleted for SMAD6 alone. At the mechanistic level, blockade of actomyosin contractility or increased PI3K signaling rescued endothelial junction defects induced by SMAD6 loss. Thus, SMAD6 normally modulates ALK1 function in endothelial cells to regulate PI3K signaling and contractility, and SMAD6 loss increases signaling through ALK1 that disrupts endothelial junctions. ALK1 loss-of-function also disrupts vascular development and function, indicating that balanced ALK1 signaling is crucial for proper vascular development and identifying ALK1 as a “Goldilocks” pathway in vascular biology regulated by SMAD6.

## INTRODUCTION

Blood vessel formation involves the expansion of a primitive vascular network of endothelial cells into different organs and tissues during embryonic life (Carmeliet, 2000). As vessels remodel and mature under the influence of environmental signals such as blood flow and tissue-specific signaling, larger arteries carry blood away from the heart and veins return blood to the heart. Extensive capillary beds form between arteries and veins, and these capillaries acquire organ-specific properties that support tissue metabolism and function (Augustin and Koh, 2017; Aird, 2007; Rafii et al., 2016). Among organ-specific developmental programs, the fetal liver is unique in that it receives oxygenated blood from the placenta via the portal vein rather than arteries. Blood flows through the liver parenchyma via capillaries that over time specialize into sinusoids comprised of liver sinusoidal endothelial cells (LSEC), then returns to the heart via the central vein (Swartley et al., 2016; Lammert et al., 2003). As LSEC differentiate they down-regulate some capillary markers and up-regulate markers not normally expressed by blood capillaries, forming a uniquely discontinuous basement membrane with fenestrations that permits filtration of blood components (Koch et al., 2021; Poisson et al., 2017). In mice, LSEC maturation starts embryonically around E12 and increases through post-natal stages (Matsumoto et al., 2001; Gómez-Salinero et al., 2022). Thus, development of the fetal liver vasculature is unique in several ways but remains poorly understood.

Among numerous regulators of vascular development, the BMP (bone morphogenetic protein) signaling pathway is essential for proper blood vessel formation in ways that are complex and context-dependent (Kulikauskas et al., 2022; García de Vinuesa et al., 2016; David et al., 2009; Moreno-Miralles et al., 2009). Canonical BMP signaling utilizes extracellular ligands that bind hetero-tetrameric complexes of Type I and Type II trans-membrane receptors. Within a complex, Type II receptors phosphorylate Type I receptors to activate Type I receptor phosphorylation of cytosolic receptor-mediated SMADs (R-SMADs). This phosphorylation allows R-SMADS to form heterotrimers with SMAD4 that translocate to the nucleus to transcriptionally regulate target genes. BMP signaling in endothelial cells leads to both pro-angiogenic vascular phenotypes such as tip cell formation and sprouting, and homeostatic vascular phenotypes associated with quiescence (Bautch, 2019; Mouillesseaux et al., 2016; Larrivée et al., 2012; Goumans et al., 2018). This apparent paradox of opposing phenotypic outputs likely results from different combinations of BMP complex components leading to differential responses to proliferation cues and blood flow, an important mechanical input for vascular BMP signaling (Kulikauskas et al., 2022; Baeyens et al., 2016). A key ligand regulating vascular homeostasis, BMP9, is synthesized in the liver and contributes to liver fibrosis in adult sinusoids downstream of LSEC de-differentiation (Bidart et al., 2012; Breitkopf-Heinlein et al., 2017). A closely related ligand, BMP10, is required for post-embryonic vascular development and maintenance in zebrafish and at some sites of mammalian angiogenesis and wounding (Capasso et al., 2020; Choi et al., 2022); however, how BMP signaling functions in developing liver blood vessels is not known.

ACVRL1/ALK1 (ALK1) is a BMP Type I receptor and the most avid binding partner of BMP9/10 (Scharpfenecker et al., 2007; Townson et al., 2012; Brown et al., 2005). ALK1 is also the most abundantly expressed BMP Type I receptor on most endothelial cells, and it functions in signaling complexes important for flow-mediated remodeling (Benn et al., 2017; Baeyens et al., 2016). Germline loss-of-function mutations in ALK1 are genetically linked with Hereditary Hemorrhagic Telangiectasia type II (HHT2), wherein patients exhibit microvascular hemorrhage and arteriovenous malformations (AVMs) with a liver tropism (Johnson et al., 1996). Murine genetic loss of *Alk1* is embryonic lethal due to hemorrhage and AVMs (Park et al., 2008b; Urness et al., 2000), and loss of endothelial *Alk1* function in postnatal mice results in AVMs, dilated vessels, and vascular hyperplasia in several vascular beds (Tual-Chalot et al., 2014; Ola et al., 2018; Lee et al., 2017), highlighting its role in vascular development and remodeling. ALK1 inhibits cell proliferation and migration of cultured endothelial cells (Lamouille et al., 2002; David et al., 2007b) downstream of BMP9/10 engagement (Li et al., 2020; Park et al., 2012), and laminar flow sensitizes endothelial cells to BMP9-mediated ALK1 signaling (Baeyens et al., 2016). Thus, endothelial cell ALK1 signaling is thought to promote vascular quiescence and vessel integrity via effects on proliferation and migration, although its role in organ-specific vascular development is not well-described.

Several cytosolic inhibitory SMADs (i-SMADs) also regulate BMP signaling. SMAD6 is a negative regulator of BMP signaling expressed in vascular endothelial cells whose expression is upregulated by laminar flow (Topper et al., 1997; Miyazawa and Miyazono, 2017; Hata et al., 1998; Ruter et al., 2021). SMAD6 expression is tightly regulated developmentally, and it is expressed first in the cardiac outflow tract and dorsal aorta at E9.5 of mouse gestation (Galvin et al., 2000). As development proceeds, mice carrying a *LacZ* reporter in the *Smad6* locus had expanded *LacZ* expression in endothelial cells of great arteries and other large arteries, and this expression remained strong into adulthood (Wylie et al., 2018; Galvin et al., 2000). We showed that global *Smad6* loss leads to vascular hemorrhage and late embryonic lethality (Wylie et al., 2018). Moreover, SMAD6 is transcriptionally regulated by Notch, and Notch-mediated Smad6 function negatively regulates endothelial cell BMP responsiveness and angiogenic sprouting (Mouillesseaux et al., 2016). SMAD6 is required for flow-mediated endothelial cell alignment downstream of Notch, and it stabilizes endothelial cell adherens junctions and promotes barrier function *in vitro* (Wylie et al., 2018; Ruter et al., 2021). Thus, SMAD6 promotes vessel stability and endothelial cell junction integrity, but its functional BMP pathway target in endothelial cells is not known.

Here we investigated how SMAD6 affects vascular development. We found that *Smad6* functions in endothelial cells to regulate vascular integrity, and that *Smad6* antagonizes *Alk1* function *in vivo* and in *vitro*. The developing fetal liver was particularly sensitive to disrupted *Alk1* signaling induced by *Smad6* loss, with extensive vessel hemorrhage that was rescued by reduced *Alk1* gene dosage. SMAD6 modulated ALK1-dependent endothelial cell contractility and PI3K signaling to regulate junction integrity and barrier function, revealing that SMAD6 contributes to balanced ALK1 signaling that is important for proper levels of PI3K activity and vessel integrity.

## RESULTS

### *Smad6* Functions in Endothelial Cells during Embryogenesis

To begin detailed investigations of *Smad6* function during mammalian development, we first generated mice carrying the *Smad6-lacZ* knock-in allele (*Smad6^tm1Glvn^*) on the C57BL6J background (N>10 backcross generations). Mice homozygous for this allele are hereafter referred to as *Smad6*^-/-^ global mutant mice. Heterozygous intercrosses (*Smad6^+/-^* x *Smad6^+/-^*) confirmed significant loss of *Smad6^-/-^* pups at P0, similar to the lethality documented on a mixed genetic background (Wylie et al, 2018), and genotype analysis of embryos (E12.5-E18.5) showed expected Mendelian ratios of *Smad6^-/-^* mutant embryos up to E16.5, with fewer mutant embryos identified on subsequent days **(Supp. Fig1A)**. Since *Smad6^-/-^* global mutant embryos had perturbed vascular development at E16.5, we performed subsequent analysis at this timepoint **(Fig. 1A)**. E16.5 *Smad6^-/-^* vascular phenotypes included abdominal and jugular (neck area) hemorrhage, and some embryos exhibited edema, paleness, and blood-filled dermal lymphatics with lower penetrance (data not shown). All vascular phenotypes in *Smad6^-/-^* global mutant embryos were variably penetrant, indicating that stochastic processes may contribute to the phenotype. Using semi-quantitative phenotype scoring of whole embryos **(Supp. Fig. 2A)**, we found that 100% of *Smad6^-/-^* global mutant embryos exhibited hemorrhage in at least one location **(Fig. 1C).** Abdominal hemorrhage was the most penetrant phenotype, with 97% of *Smad6^-/-^* global mutant embryos affected, while 48% of E16.5 global *Smad6^-/-^* mutant embryos had jugular hemorrhage **(Fig. 1D-E).**

**Figure 1.**
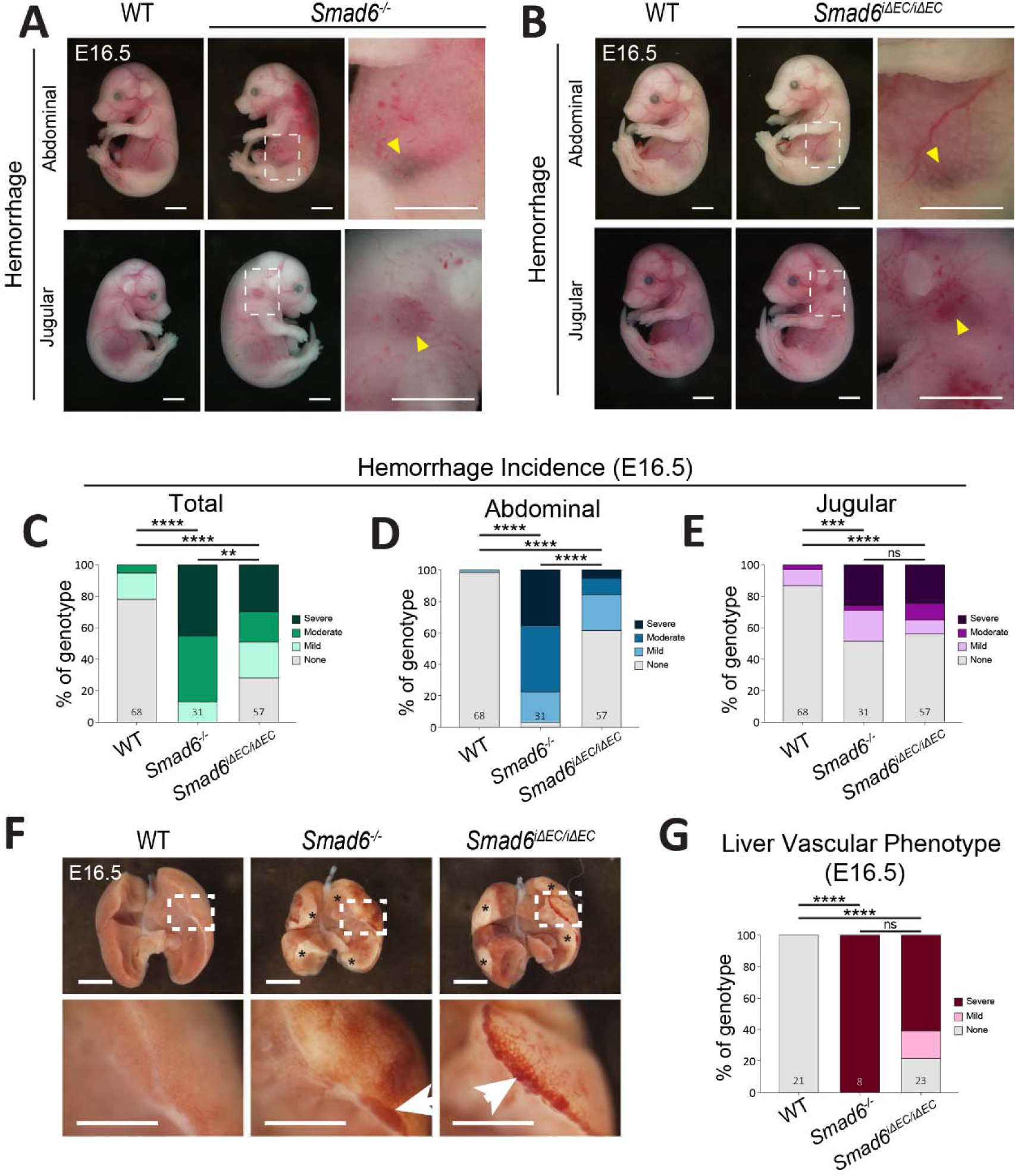
*Smad6* functions in embryonic endothelial cells. **(A, B)** Representative images of E16.5 *Smad6^-/-^* **(A)** or *Smad6^i^*^Δ^*^EC/i^*^Δ^*^EC^* **(B)** embryos and control littermates. Arrowheads, jugular or abdominal hemorrhage. Scale bar, 2.5mm. **(C-E)** Semi-quantitative E16.5 whole embryo hemorrhage analysis of indicated types (see **Supp. Fig 2A** for scoring criteria). WT, n=68; *Smad6^-/-^*, n = 31; *Smad6^i^*^Δ^*^EC/i^*^Δ^*^EC^*, n=57 embryos. **(F)** Representative images of E16.5 *Smad6^-/-^* and *Smad6^i^*^Δ^*^EC/i^*^Δ^*^EC^* livers and controls. Asterisks, pale regions. Arrows, regions of pooled blood/hemorrhage. Scale bar, 1mm. **(G)** Semi-quantitative E16.5 whole liver hemorrhage/dilation phenotype analysis (see **Supp. Fig. 2B** for scoring criteria). WT, n=21; *Smad6^-/-^,* n=8; *Smad6^i^*^Δ^*^EC/i^*^Δ^*^EC^*, n=23 livers. **, P<0.01; ***, P<0.001; ****, P<0.0001; ns, not significant. Statistics, Χ^2^ analysis.

**Figure 2.**
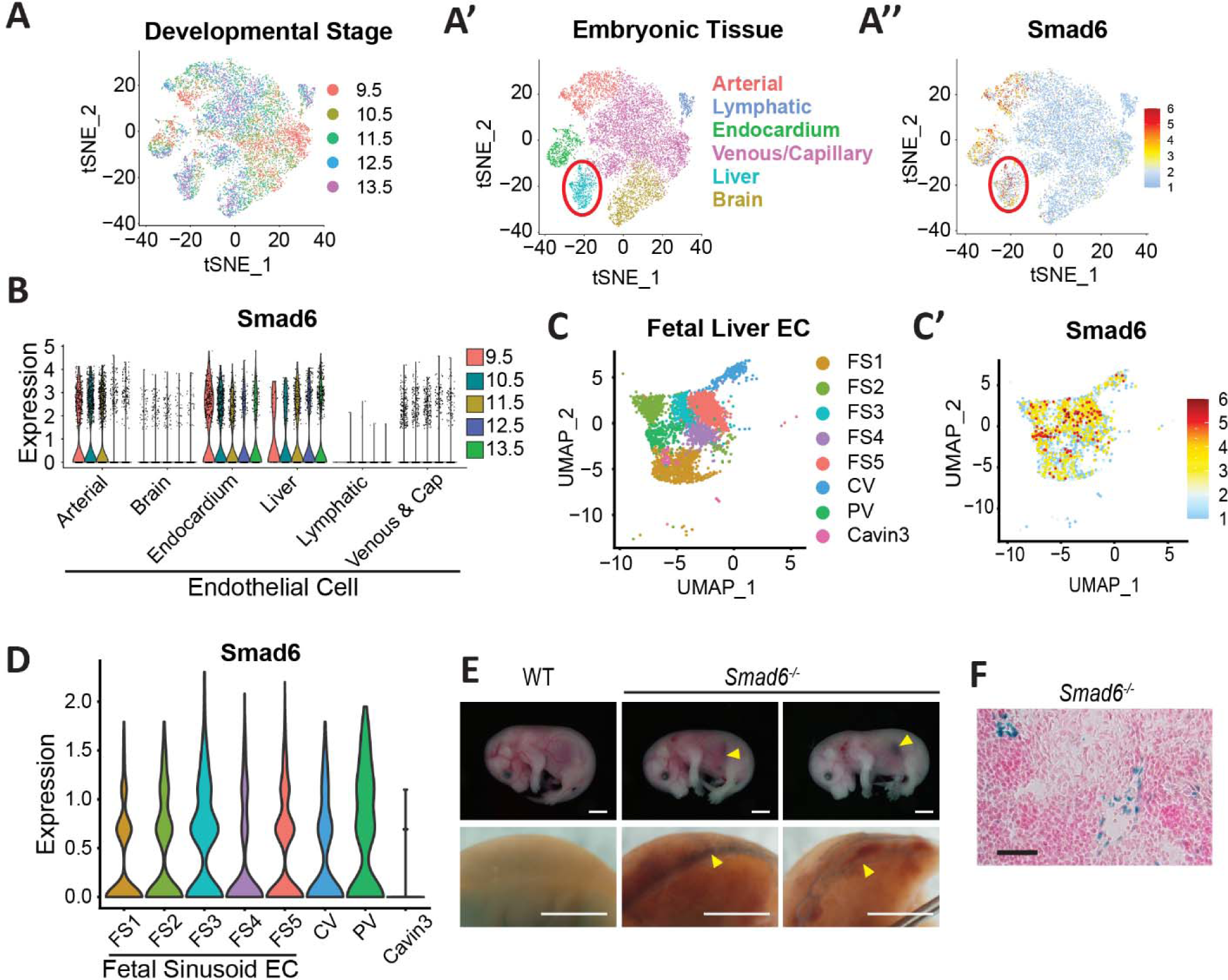
Smad6 is expressed in embryonic liver endothelial cells. **(A-B)** Re-analysis of MOCA scRNA-seq dataset. tSNE plots label endothelial cell populations by embryonic stage **(A)**, embryonic tissue **(A’)**, and Smad6 expression **(A’’)**. Red circled area denotes embryonic liver endothelial cells. Violin plot showing Smad6 expression in endothelial cell clusters by stage **(B)**. **(C-D)** Re-analysis of scRNA-seq data from (Gómez-Salinero et al., 2022). UMAP plot of fetal liver endothelial cell clusters by stage (E12-E18) **(C)** and Smad6 expression **(C’). (D)** Violin plot of Smad6 expression levels in fetal liver endothelial cell clusters. FS1-5, fetal sinusoidal endothelial cell populations; CV, central vein; PV, portal vein; Cavin3, hepatic artery marker. **(E)** (Top) E16.5 *Smad6^-/-^* global mutant embryos and control. Arrowheads, abdominal hemorrhage. Scale bar, 2.5mm. (Bottom) Isolated livers from same embryos wholemount stained for LacZ. Arrowheads, liver vessel dilation/hemorrhage. Scale bar, 1mm. **(F)** E16.5 liver cryosection from *Smad6^-/-^* global mutant embryo, stained for *lacZ* and counterstained for nuclear fast red. Scale bar, 100µm.

We next asked whether *Smad6* function is required cell-autonomously in vascular endothelial cells during mammalian development. A *Smad6* floxed allele was generated by inserting LoxP sites around exon 4, which encodes a MH2 protein domain required for function (**Supp. Fig. 1B**). Mice carrying this allele were bred to an endothelial cell-specific tamoxifen-inducible line, *Tg(Cdh5-cre/ERT2)1Rha* (hereafter referred to as *Cdh5-Cre^ERT2^*), and excision was induced at E10.5 via tamoxifen oral gavage (**Supp. Fig. 1C**). We attempted to analyze pups at birth, but they required harvest via C-section just prior to birth due to maternal distress. None of the 4 pups genotyped as *Smad6^fl/fl^;Cdh5-Cre^ERT2^* (*Smad6^i^*^Δ^*^EC/i^*^Δ^*^EC^*) were viable at E20.5 harvest, while 7 non-mutant pups were alive at this time (**Supp. Fig. 1D**); these findings indicate that *Smad6^i^*^Δ^*^EC/i^*^Δ^*^EC^* embryos did not survive to birth, similar to *Smad6*^-/-^ global mutant embryos. Phenotype scoring of whole *Smad6^i^*^Δ^*^EC/i^*^Δ^*^EC^* mutant embryos at E16.5 recapitulated *Smad6^-/-^* global mutant vascular phenotypes, with significant abdominal and jugular hemorrhage compared to littermate controls (**Fig. 1B-E**), and sporadic edema, paleness, and blood-filled lymphatics (data not shown). These findings indicate that the primary vascular phenotypes associated with *Smad6* global loss are largely endothelial cell-specific and show that *Smad6* function is required in endothelial cells for vascular integrity *in vivo*.

The prevalence of abdominal hemorrhage suggested liver involvement, and examination of isolated E16.5 livers revealed significant vessel dilation and hemorrhage in both *Smad6^-/-^* global and *Smad6^i^*^Δ^*^EC/i^*^Δ^*^EC^* livers, with pale regions not seen in controls (**Fig. 1F-G**). The phenotype scoring of isolated livers (**Supp. Fig2B**) was similarly penetrant between the classes of mutant embryos, suggesting that whole embryo abdominal scoring was less sensitive and that *Smad6* functions in embryonic liver endothelial cells.

We hypothesized that the residual reduced penetrance of vascular phenotypes in *Smad6^i^*^Δ^*^EC/i^*^Δ^*^EC^* embryos compared to *Smad6^-/-^* global mutants might reflect later or less complete removal of *Smad6* from endothelial cells. We used a tamoxifen-inducible global Cre line, *UBC-Cre^ERT2^*, to generate *Smad6^i^*^Δ^*^/i^*^Δ^ E16.5 embryos using the same excision protocol as for *Smad6*^Δ^*^ECi^*^Δ^*^EC^* embryos and found that the severity of vascular phenotypes mirrored that of the *Smad6^-/-^* global mutants (**Supp. Fig. 3A-G**). Thus, *Smad6* deletion mid-gestation did not lead to significant reduced phenotype penetrance, and the *Smad6^-/-^* and *Smad6^i^*^Δ^*^/i^*^Δ^ lines were used interchangeably in subsequent experiments. For *Smad6^i^*^Δ^*^EC/i^*^Δ^*^EC^* embryos, gene excision via PCR analysis of lung lysates at E16.5 revealed a predicted excision band not seen in controls (**Supp. Fig. 3H-I**). We next analyzed Smad6 liver expression in E16.5 *Smad6^i^*^Δ^*^EC/i^*^Δ^*^EC^* embryos and found that isolated PECAM1-enriched cell populations had significantly reduced levels of Smad6 RNA that correlated with phenotype severity (**Supp Fig 3 J-K**), indicating that embryo-dependent partially inefficient excision may be responsible for residual reduced penetrance of the liver vascular phenotype in *Smad6^i^*^Δ^*^EC/i^*^Δ^*^EC^* embryos.

**Figure 3.**
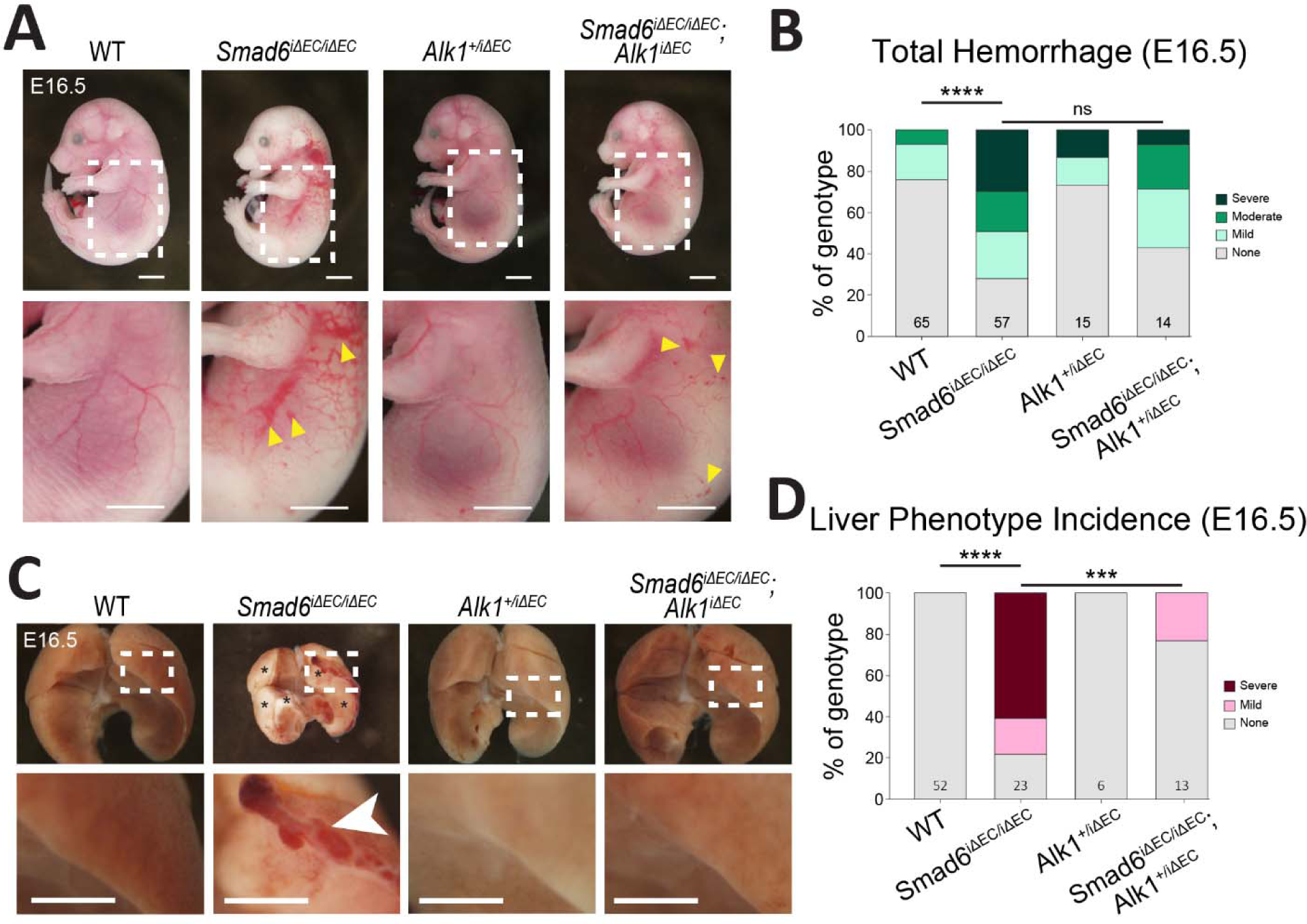
Reduced *Alk1* gene dosage partially rescues *Smad6i*Δ*EC/i*Δ*EC* vascular defects. **(A)** Representative images of E16.5 embryos if indicated genotypes. Arrowheads, blood in lymphatics. Scale bar, 2.5mm. **(B)** Semi-quantitative whole embryo hemorrhage analysis (see **Supp. Fig 2A** for scoring criteria). WT, n=65; *Smad6^i^*^Δ^*^EC/i^*^Δ^*^EC^*, n=57; *Alk1^+/i^*^Δ^*^EC^*, n=15; *Smad6^i^*^Δ^*^EC/i^*^Δ^*^EC^*;*Alk^+/i^*^Δ^*^EC^*, n=14 embryos. **(C)** Representative images of E16.5 livers of indicated genotypes. Asterisks, pale regions. Arrows, regions of pooled blood. Scale bar, 1mm. **(D)** Semi-quantitative E16.5 whole liver phenotype analysis (see **Supp. Fig. 2B** for scoring criteria). WT, n=52; *Smad6^i^*^Δ^*^EC/i^*^Δ^*^EC^*, n=23; *Alk^+/i^*^Δ^*^EC^,* n=6; *Smad6^i^*^Δ^*^EC/i^*^Δ^*^EC^*;*Alk^+/i^*^Δ^*^EC^,* n=13 livers. ***, P<0.001; ****, P<0.0001. Statistics, Χ^2^ analysis.

### *Smad6* is Expressed in Embryonic Liver Endothelial Cells

The hepatic vascular defects seen in *Smad6^i^*^Δ^*^EC/i^*^Δ^*^EC^* embryos were interesting, as the embryonic liver vasculature consists of veins and capillaries at this stage, and hepatic artery formation is only detectable just before birth (Swartley et al., 2016). In contrast, robust embryonic *Smad6* expression via the *lacZ* reporter readout was documented in larger arteries and the outflow tract that did not have obvious defects (data not shown) (Wylie et al., 2018; Galvin et al., 2000). To more rigorously examine vascular Smad6 expression, we reanalyzed several published single-cell (sc) RNA seq datasets. scRNA seq data from an EC Atlas of adult mouse tissues (Kalucka et al., 2020) revealed *Smad6* expression in endothelial cells of several organs **(Supp. Fig 4A-A’)**; *Smad6* expression was substantial in vein and capillary endothelial cells along with arterial expression in liver and lung, while expression was more localized to arterial endothelial cells in the brain **(Supp. Fig 4B-D, B’-D’)**. Re-analysis of a second dataset from adult mouse brain and lung endothelial cells (Vanlandewijck et al., 2018; He et al., 2018) revealed *Smad6* RNA expression in venous and capillary endothelial cells, albeit at a lower prevalence than in arterial cells **(Supp. Fig 4E-F)**. Additionally, Gomez-Salinero et al. re-analyzed the Tabula Muris database for highly expressed liver genes and identified Smad6 expression as enriched in liver endothelial cells (Gómez-Salinero et al., 2022). Taken together, these data indicate that Smad6 is expressed in endothelial cells from all caliber adult mammalian blood vessels, including veins and capillaries.

**Figure 4.**
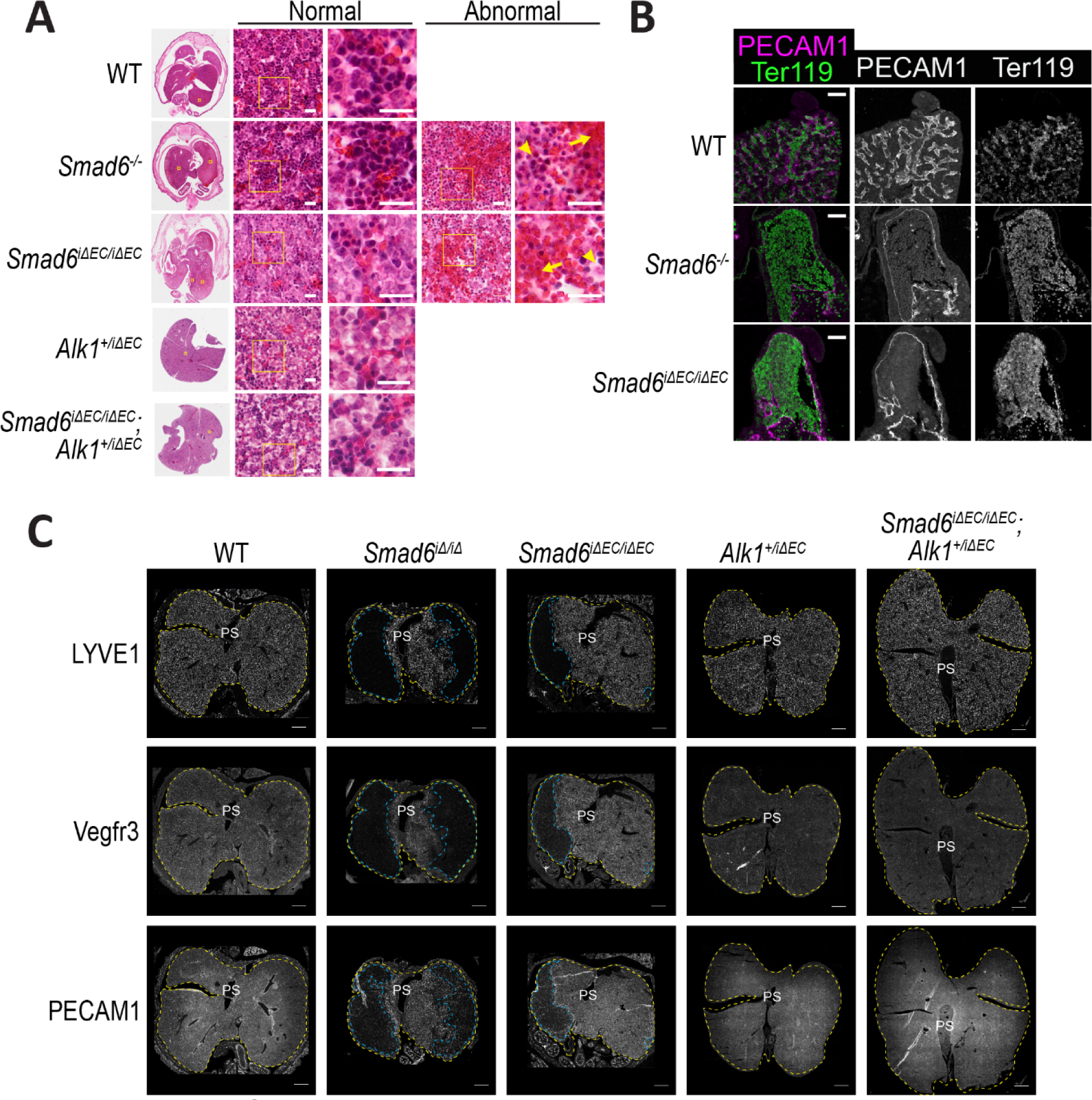
Endothelial *Smad6* maintains embryonic liver vessels via *Alk1* regulation. **(A-C)** Representative images of E16.5 liver sections of indicated genotypes. **(A)** H&E stain. Far left, whole livers. Yellow outlined areas are magnified to right. Middle, areas of normal liver parenchyma in livers of indicated genotypes. Yellow outlined areas to left are magnified to right. Far right, areas of abnormal parenchyma in *Smad6* mutant liver sections. Yellow outlined areas to left are magnified to right. Scale bar, 20µm. Arrow, hemorrhage; arrowhead, tissue disorganization. **(B)** Representative immunofluorescence images for PECAM1 (endothelial cells) and Ter119 (red blood cells) at periphery of E 16.5 livers of indicated genotypes. Scale bar, 50µm. **(C)** Representative images of immunofluorescence of E16.5 liver sections (adjacent sections for each liver) for indicated liver endothelial markers. Yellow dashed line, liver outline; blue dashed line, avascular areas. Scale bar, 500µm.

To assess embryonic *Smad6* endothelial cell expression, we re-analyzed the Mouse Organogenesis Cell Atlas (MOCA) dataset (Cao et al., 2019) by extracting endothelial cell information by embryonic stage (E9.5-13.5) and organ **(Fig. 2A-A’)**. This analysis revealed substantial Smad6 expression in embryonic (E9.5-13.5) arterial, endocardial, and liver endothelial cells **(Fig. 2A’’, B)**. Next, a recently published scRNA seq dataset of mouse liver endothelial cells from E12-P30 (Gómez-Salinero et al., 2022) was re-analyzed with a focus on embryonic stages, revealing Smad6 expression in most liver embryonic endothelial cell clusters from E12-E18, including expression in the larger portal and central veins and fetal sinusoidal endothelial cells (FS1-FS5) **(Fig. 2C-C’, D)**. Finally, we assessed Smad6 expression in E16.5 embryonic livers via the *lacZ* reporter and documented *lacZ* expression in surface vessels and in endothelial cells lining hepatic vessels **(Fig. 2E-F)**, consistent with the scRNA seq data and with the conclusion that Smad6 is expressed in veins and capillaries of the embryonic liver.

### *Smad6* Exhibits Epistasis with *Alk1* in Embryonic Liver Blood Vessels

Having established that endothelial cell Smad6 function regulates vessel integrity in the embryonic liver, we next asked how SMAD6 exerts its effects, and we hypothesized that some aspect of BMP function is normally regulated by SMAD6. We turned our attention to the BMP Type I receptor Alk1, as signaling through complexes containing this receptor is important for vascular integrity and flow-mediated responses (Baeyens et al., 2016; Tual-Chalot et al., 2014). We hypothesized that SMAD6 antagonizes ALK1 signaling and predicted that genetic reduction of *Alk1* would rescue the loss of vascular integrity seen with *Smad6* loss. Global *Alk1* deletion is lethal at mid-gestation with impaired vascular development and vessel dilation (Oh et al., 2000), and homozygous endothelial cell deletion of *Alk1* is lethal within several days of excision in neonates due to AVMs and pulmonary hemorrhage (Tual-Chalot et al., 2014; Park et al., 2008b). We confirmed that endothelial-specific deletion of *Alk1* starting at E10.5 was embryonic lethal at E16.5 and likely earlier, since the mutant embryos were partially resorbed at this time point **(Supp. Fig 5A)**. We showed that concomitant deletion of *Smad6* did not alter *Alk1*-dependent lethality with this excision protocol **(Supp. Fig 5B)**, so we tested genetic epistasis in embryos with one *Alk1* allele and both *Smad6* alleles deleted in endothelial cells. Whole embryo examination revealed a trend for increased rescue of total hemorrhage in *Smad6^i^*^Δ^*^EC/i^*^Δ^*^EC^*;*Alk^+/i^*^Δ^*^EC^* embryos relative to *Smad6^i^*^Δ^*^EC/i^*^Δ^*^EC^* embryos **(Fig. 3A-B)**, while isolated liver analysis showed highly significant rescue of liver vascular defects in embryos with reduced *Alk1* gene dosage **(Fig. 3C-D)**. None of the *Smad6^i^*^Δ^*^EC/i^*^Δ^*^EC^*;*Alk^+/i^*^Δ^*^EC^* livers presented with a severe vascular phenotype, and 77% of embryonic livers with reduced *Alk1* gene dosage had no discernable vascular defect. These results indicate that *Smad6* and *Alk1* have an epistatic relationship in embryonic liver endothelial cells and suggest that SMAD6 normally restricts Alk1 activity.

**Figure 5.**
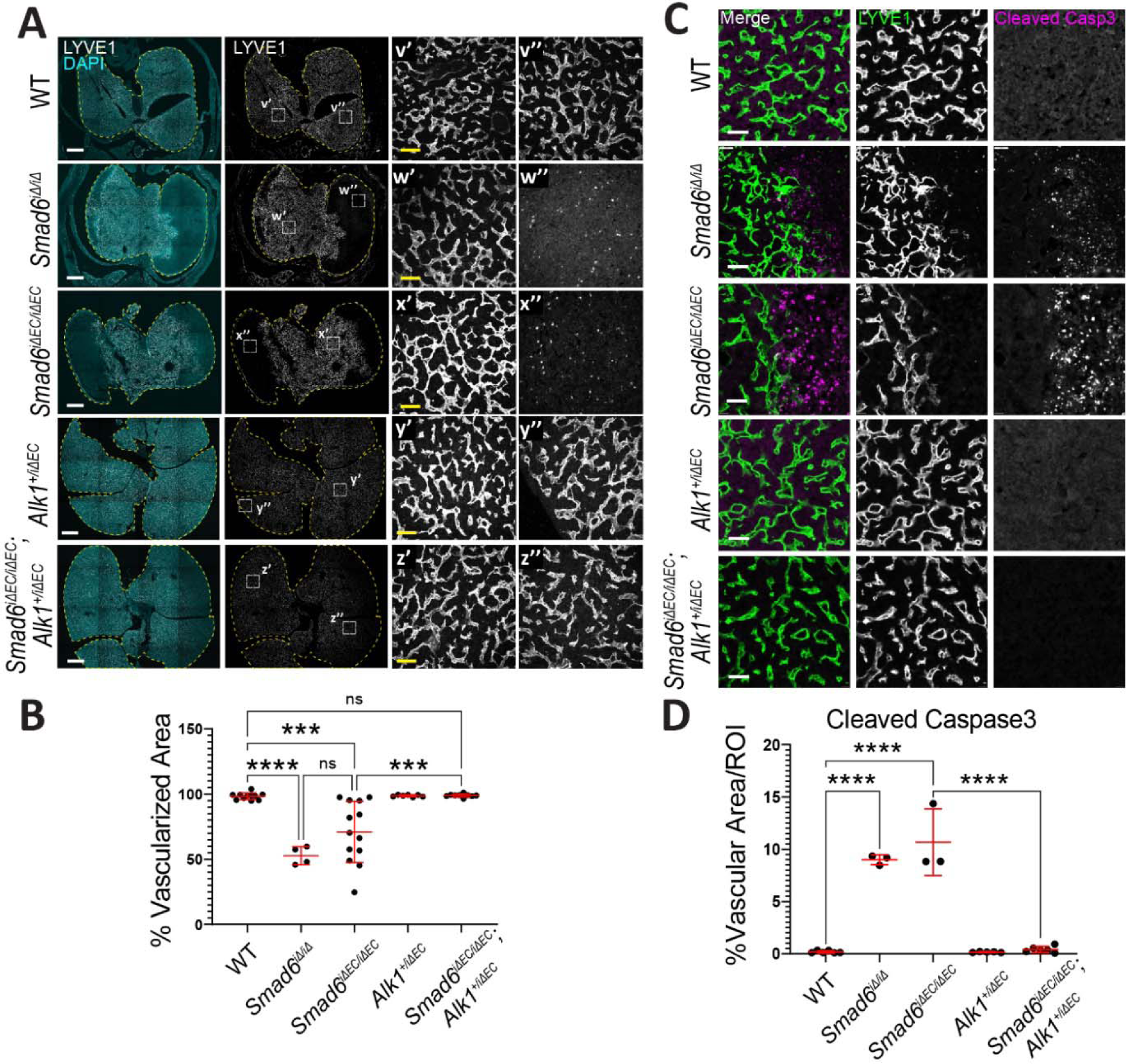
Endothelial *Smad6* maintains embryonic liver vascular patterning via *Alk1* regulation. **(A, C)** Representative images of E16.5 liver sections of indicated genotypes. **(A)** Immunofluorescence for DAPI (nucleus) and LYVE1 (liver endothelial cell). Yellow dashed line, liver outline; white dashed square, areas magnified to right. Scale bar, 500µm. Insets, LYVE1 staining in vascular areas (’) and second areas that are avascular in mutants (’’). Scale bar, 50µm. **(B)** Quantification of % LYVE1^+^ vascularized area/whole liver area. WT, n=11; *Smad6^i^*^Δ^*^/i^*^Δ^, n=4; *Smad6^i^*^Δ^*^EC/i^*^Δ^*^EC^*, n=13; *Alk1^+/i^*^Δ^*^EC^,* n=7; *Smad6^i^*^Δ^*^EC/i^*^Δ^*^EC^*;*Alk^+/i^*^Δ^*^EC^,* n=9 livers. **(C)** Representative images of immunofluorescence for LVYE1 and cleaved Caspase3. Scale bar, 50µm. **(D)** Quantification of whole liver scans of cleaved Caspase3 channel/total ROI area (region of interest). WT, n=7; *Smad6^i^*^Δ^*^/i^*^Δ^, n=3; *Smad6^i^*^Δ^*^EC/i^*^Δ^*^EC^*, n=6; *Alk1^+/i^*^Δ^*^EC^,* n=5; *Smad6^i^*^Δ^*^EC/i^*^Δ^*^EC^*;*Alk^+/i^*^Δ^*^EC^,* n=6 livers. ***, P<0.001; ****, P<0.0001; ns, not significant. Data are individual points ±SD. Statistics, one-way ANOVA with Tukey’s test to correct for multiple comparisons.

### *Smad6* Maintains Vessel Integrity in the Embryonic Liver via *Alk1* Regulation

To further understand *Smad6* function and epistasis with *Alk1* in embryonic liver vessels, we performed histological and marker analysis on embryonic liver sections. In general, microscopic phenotypes were shared by livers from *Smad6^-/-^* global mutant embryos, *Smad6^i^*^Δ^*^/i^*^Δ^ mutant embryos, and *Smad6^i^*^Δ^*^EC/i^*^Δ^*^EC^* mutant embryos, suggesting that effects of Smad6 loss on liver development result from endothelial cell-selective functions of *Smad6*. H&E stained E16.5 liver sections revealed areas of hemorrhage and loss of tissue organization in both *Smad6^-/-^* And *Smad6^i^*^Δ^*^EC/i^*^Δ^*^EC^* mutants not seen in controls **(Fig. 4A),** amongst areas that appeared relatively normal. This mosaicism was not observed in either the *Alk1^+/i^*^Δ^*^EC^* or *Smad6^i^*^Δ^*^EC/i^*^Δ^*^EC^*;*Alk^+/i^*^Δ^*^EC^* liver sections that appeared similar to controls, indicating that reduced gene dosage of *Alk1* rescued the regions of hemorrhage and disorganization. The areas of hepatic vascular loss appeared skewed towards peripheral areas of the liver lobes, and staining for the vascular marker PECAM1 (CD31) and the red blood cell marker Ter119 revealed large and dilated peripheral vessels in *Smad6^-/-^* And *Smad6^i^*^Δ^*^EC/i^*^Δ^*^EC^* mutant livers (**Fig. 4B**), consistent with the whole liver phenotype.

Staining of adjacent sections for Lyve1 (marker of liver sinusoidal endothelial cells (LSEC)), Vegfr3 (early endothelial cell marker) or PECAM1 (pan-endothelial cell marker) revealed co-incident expression or loss of expression in liver capillaries of *Smad6* mutant embryos, indicating avascular areas within the liver parenchyma; these avascular areas were not seen in livers of *Smad6^i^*^Δ^*^EC/i^*^Δ^*^EC^*;*Alk^+/i^*^Δ^*^EC^* embryos with reduced *Alk1* gene dosage **(Fig. 4C)**. Quantification revealed that Lyve1+ area was significantly reduced in both *Smad6^-/-^* global mutant and *Smad6^i^*^Δ^*^EC/i^*^Δ^*^EC^* livers, but not in *Smad6^i^*^Δ^*^EC/i^*^Δ^*^EC^*;*Alk^+/i^*^Δ^*^EC^* livers with reduced *Alk1* dosage **(Fig. 5A-B)**, confirming that peripheral mosaic avascularity in the liver parenchyma is a hallmark of endothelial *Smad6* loss and that reduced *Alk1* gene dosage rescues this phenotype.

Examination of avascular areas in the mutant livers revealed significant levels of cleaved caspase3 in regions that border vascularized areas, indicative of parenchymal cell death that was not seen in controls or *Smad6^i^*^Δ^*^EC/i^*^Δ^*^EC^*;*Alk^+/i^*^Δ^*^EC^* mutant livers with reduced *Alk1* gene dosage **(Fig 5C-D)**. Thus, endothelial loss of Smad6 function leads to mosaic loss of capillary vessels in the embryonic liver parenchyma accompanied by cell death that is dependent on *Alk1* gene dosage, indicating that *Smad6* restriction of *Alk1* signaling normally regulates vascular integrity during liver embryogenesis.

### *Smad6* Promotes Liver Sinusoidal Endothelial Cell Differentiation

Liver sinusoidal endothelial cells (LSEC) are specialized liver capillary vessels with a discontinuous basement membrane and fenestrations. During mid-gestation LSEC begin to form and express markers of classic blood endothelial cells, but over time they acquire LSEC-specific markers and develop a unique phenotypic and functional profile (Geraud et al., 2017). Classic endothelial cell markers such as PECAM1 and CD34 are down-regulated while LSEC markers such as Lyve1 and Stabilin-2 are upregulated (Sugiyama et al., 2010; Poisson et al., 2017; Schledzewski et al., 2011). Reversal of LSEC differentiation leads to loss of fenestration and basement membrane deposition, a process called capillarization, which leads to liver fibrosis and dysfunction in adult livers (Schaffner and Poper, 1963; Xie et al., 2012; DeLeve, 2015). We asked whether the remaining capillary beds in *Smad6* mutant livers exhibited abnormal LSEC differentiation. We found that both *Smad6^i^*^Δ^*^/i^*^Δ^ and *Smad6^i^*^Δ^*^EC/i^*^Δ^*^EC^* mutant E16.5 livers had excessive Collagen IV deposition in capillary beds that was not seen in controls or in mutant livers with reduced *Alk1* gene dosage, accompanied by ectopic staining for the smooth muscle marker αSMA **(Fig 6A-B)**. This excess of basement membrane protein indicates that LSEC of *Smad6* mutant livers are more capillarized. Further analysis revealed that the diameter of capillary vessels was significantly increased in both the *Smad6^i^*^Δ^*^/i^*^Δ^ and *Smad6^i^*^Δ^*^EC/i^*^Δ^*^EC^* E16.5 mutant livers, and this phenotype was partially rescued by reduced gene dosage of *Alk1* **(Fig. 6C)**.

**Figure 6.**
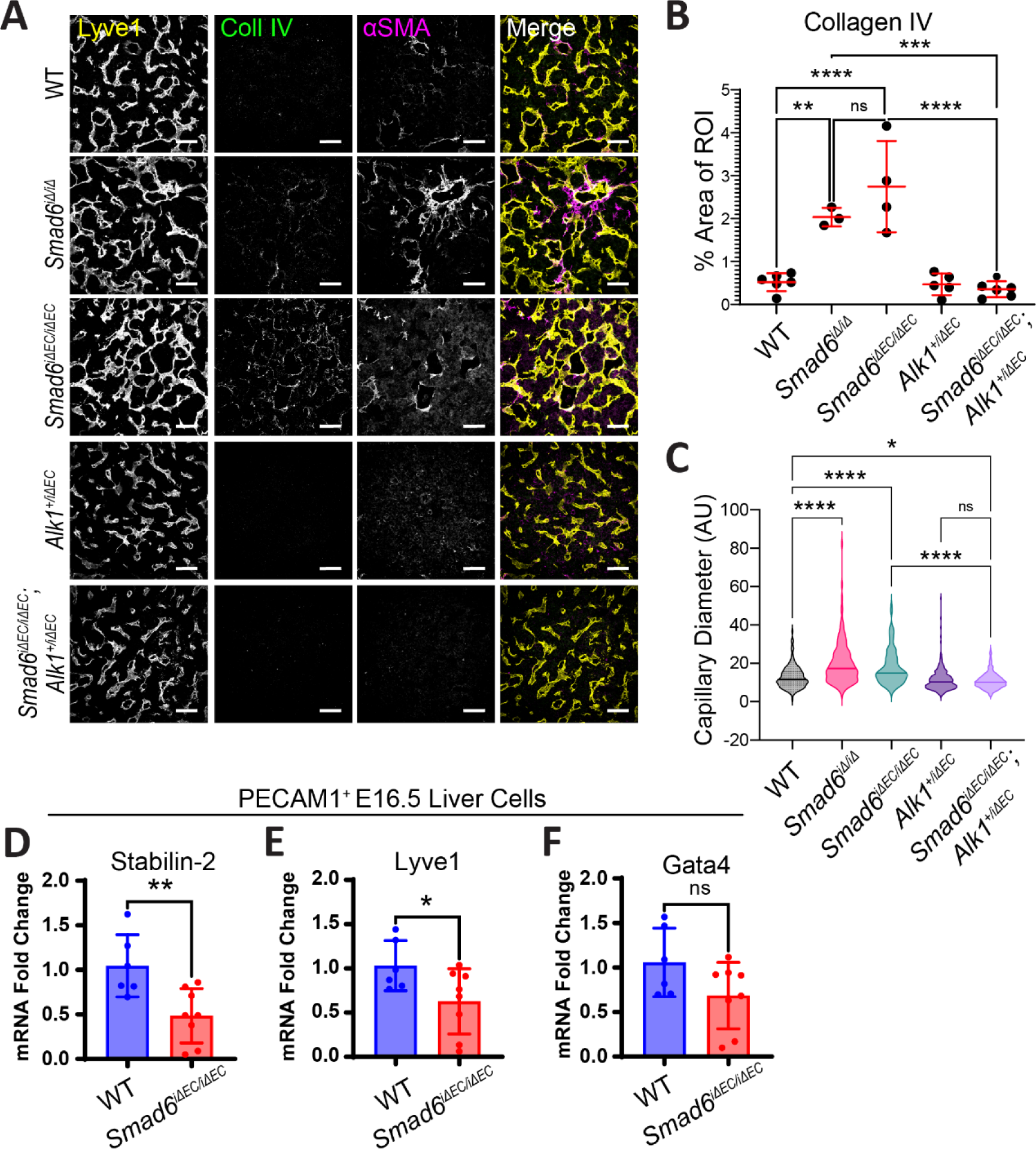
Hepatic endothelial cell *Smad6* loss augments LSEC capillarization. **(A)** Representative images of immunofluorescence staining of E16.5 liver sections of indicated genotypes for Lyve1, collagen IV and αSMA (smooth muscle actin). Scale bar, 50µm. **(B)** Collagen IV area/ROI quantification. WT, n=6; *Smad6^i^*^Δ^*^/i^*^Δ^, n=3; *Smad6^i^*^Δ^*^EC/i^*^Δ^*^EC^*, n=4; *Alk1^+/i^*^Δ^*^EC^,* n=5; *Smad6^i^*^Δ^*^EC/i^*^Δ^*^EC^*;*Alk^+/i^*^Δ^*^EC^,* n=6 livers. **(C)** Quantification of capillary diameter. WT, n=3; *Smad6^i^*^Δ^*^/i^*^Δ^, n=3; *Smad6^i^*^Δ^*^EC/i^*^Δ^*^EC^*, n=3; *Alk1^+/i^*^Δ^*^EC^,* n=5; *Smad6^i^*^Δ^*^EC/i^*^Δ^*^EC^*;*Alk^+/i^*^Δ^*^EC^,* n=6 livers. **(D-F)** qRT-PCR of PECAM1-enriched cells from E16.5 *Smad6^i^*^Δ^*^EC/i^*^Δ^*^EC^* and control littermate livers for **(D)** Stabilin-2, **(E)** Lyve1, and **(F)** Gata4. CT values normalized to Gapdh and mRNA expression reported as fold change relative to WT average. WT, n=6; *Smad6^i^*^Δ^*^EC/i^*^Δ^*^EC^*, n=8 livers. *, P<0.05; **, P<0.01. Data, individual points for each embryo ±SD. Statistics, one-way ANOVA with Tukey’s test to correct for multiple comparisons.

To better define the differentiation status of LSEC in embryonic mutant livers lacking endothelial *Smad6* function, we enriched for PECAM1^+^ endothelial cells from dissociated E16.5 livers and performed qRT-PCR for expression of relevant genes. RNA levels of the LSEC maturation markers, Stabilin-2 and Lyve1, were significantly decreased in *Smad6^i^*^Δ^*^EC^*^/^*^i^*^Δ^*^EC^* liver endothelial cells compared to controls **(Fig. 6D, E)**, consistent with the idea that liver vessels lacking *Smad6* are more capillarized than stage-matched controls. Expression of Gata4, a transcription factor that regulates LSEC differentiation (Geraud et al., 2017), trended down in *Smad6^i^*^Δ^*^EC/i^*^Δ^*^EC^* liver endothelial cells although it did not reach significance **(Fig. 6F)**. The PECAM-negative fraction of dissociated livers showed no significant changes in vascular markers **(Supp. Fig 6A-C).**

### SMAD6 Antagonizes ALK1 in Endothelial Cells to Maintain Junction Integrity and Flow Responses

SMAD6 stabilizes endothelial adherens junctions, maintains vascular barrier function, and is required for flow-mediated endothelial cell alignment (Wylie et al., 2018; Ruter et al., 2021). To define the cellular mechanism of SMAD6 function and further explore its relationship to ALK1 signaling in endothelial cells, we depleted RNA levels in primary human endothelial cells (HUVEC) using siRNA knockdown. Absent flow, we confirmed that endothelial cells depleted for Smad6 have adherens junctions that appear destabilized compared to controls. In contrast, endothelial cells depleted for Alk1 have linear junctions that appear stable, and concurrent depletion of Smad6 and Alk1 rescues the destabilized junction morphology seen with Smad6 depletion **(Fig. 7A, insets**). Under laminar flow conditions these relationships remained, and concurrent depletion of Smad6 and Alk1 rescued the mis-alignment seen with Smad6 depletion alone **(Fig. 7A-B)**. We confirmed that endothelial cell junction de-stabilization was phenocopied by addition of BMP9, and junction morphology was normalized with Alk1 depletion with or without Smad6 depletion upon BMP9 exposure **(Supp. Fig. 7)**, indicating that the effects of Alk1 on junctions are downstream of BMP9. We next functionally assessed monolayer integrity using an adapted protocol (Dubrovskyi et al., 2013) that reveals biotin-labeled matrix accessible to streptavidin, and found that labeling was significantly increased over controls in endothelial cells depleted for Smad6 under both static and flow conditions, and rescued to control levels with concurrent depletion of Smad6 and Alk1 **(Fig. 7C-E)**. We further analyzed functional effects of Smad6 and Alk1 on endothelial junctions by measuring electrical resistance across monolayers using Real Time Cell Analysis (RTCA). We confirmed that cells depleted for Smad6 had reduced electrical resistance compared to controls, and this disruption was rescued by concurrent Alk1 depletion **(Fig. 7F)**. These findings are consistent with the idea that SMAD6 regulates endothelial junctions and manages flow responses via negative modulation of BMP9/ALK1-dependent signaling.

**Figure 7.**
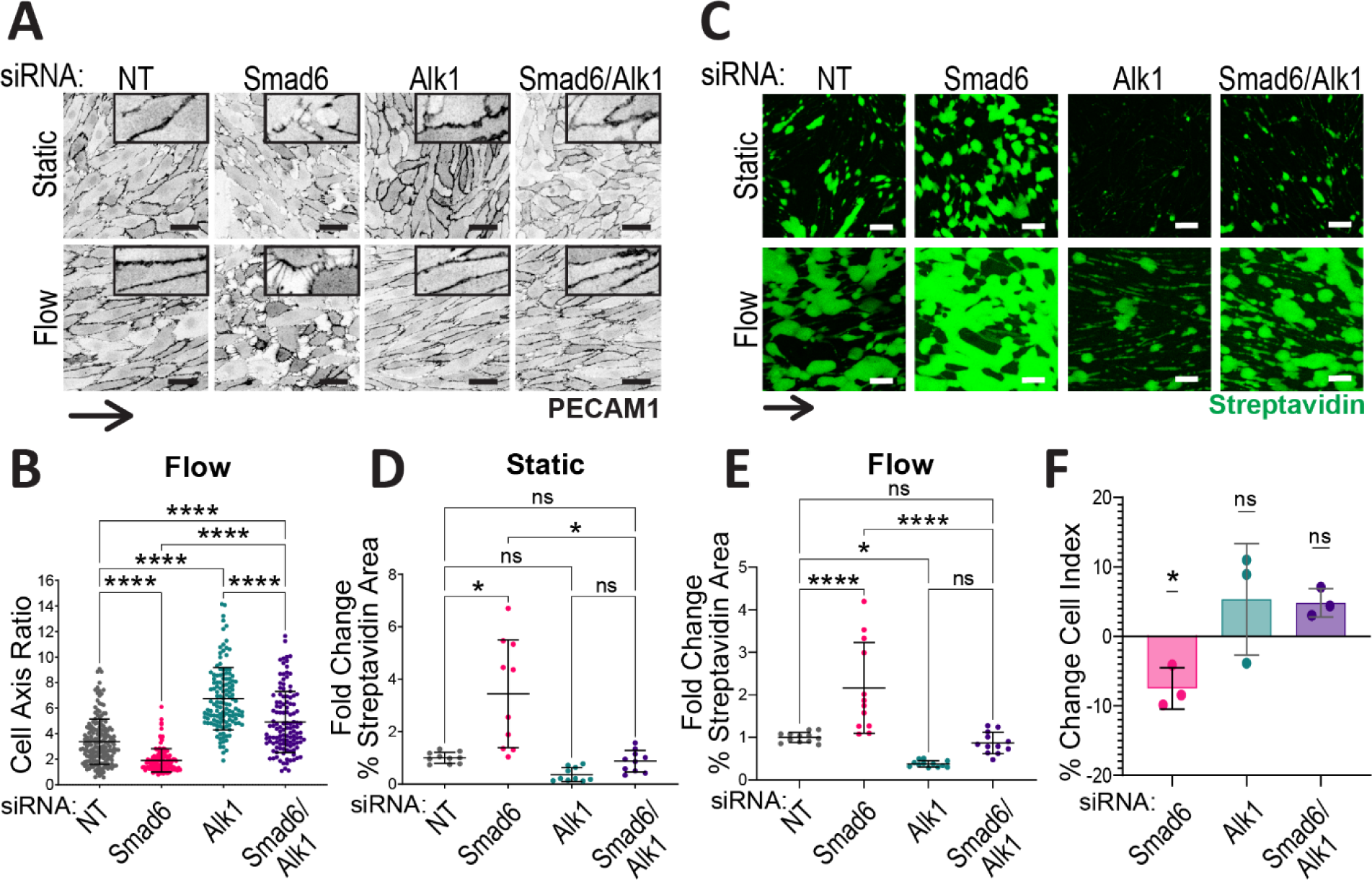
SMAD6 interacts with ALK1 in endothelial cells to maintain junction integrity and flow responses. HUVEC treated with non-targeting (NT), Smad6, and/or Alk1 siRNA were exposed to static conditions or laminar flow (7.5 dyn/cm^2^/72hr). **(A)** Cells stained for PECAM1. Scale bar, 50µm. **(B)** Quantification of cell axis ratio under flow. Data are mean ± SD (each data point = 1 cell), n ≥ 3 replicates per condition. **(C)** Cells cultured on biotinylated fibronectin under static or flow conditions and incubated with Streptavidin-488. Scale bar, 50µm. **(D-E)** Quantification of % streptavidin-488 area/total FOV (field of view) under static **(D)** or flow conditions **(E)**. Data are mean ± SD (each data point = 1 field of view), with n ≥ 3 experimental replicates per condition. *, P<0.05; **, P<0.01; ****, P<0.0001; ns, not significant. Statistics, one-way ANOVA with Tukey’s test to correct for multiple comparisons. **(F)** Quantification of % change in cell index for RTCA measured at 24hr. Normalized to NT siNRA control cell index. Data are mean ± SD (each data point an experimental replicate), with n = 3 replicates per condition. *, P<0.05; ns, not significant. Statistics, one sample t-test comparing conditions to hypothetical mean of 0.

### SMAD6 Regulates Endothelial Cell Contractility and PI3K Signaling via ALK1

The destabilized junction morphology of EC depleted for Smad6 was reminiscent of hypercontractility, so we hypothesized that SMAD6 regulates endothelial cell contractility. The contractility agonist thrombin induced junction destabilization that phenocopied the junction morphology induced by Smad6 depletion in endothelial cells, while contractility blockade via blebbistatin led to a more linear junction morphology in Smad6 silenced endothelial cells, indicating that Smad6 depletion regulates endothelial cell contractility **(Fig. 8A)**. Alk1 depletion blunted thrombin-induced junction destabilization independent of Smad6 depletion. Functionally, thrombin treatment significantly increased biotin matrix labeling in all conditions over similarly depleted controls, while contractility blockade rescued the increased matrix labeling seen with Smad6 depletion **(Fig. 8B-C)**. These results indicate that SMAD6 is required to modulate and prevent endothelial cell hypercontractility, and that this effect goes through ALK1 signaling.

**Fig. 8.**
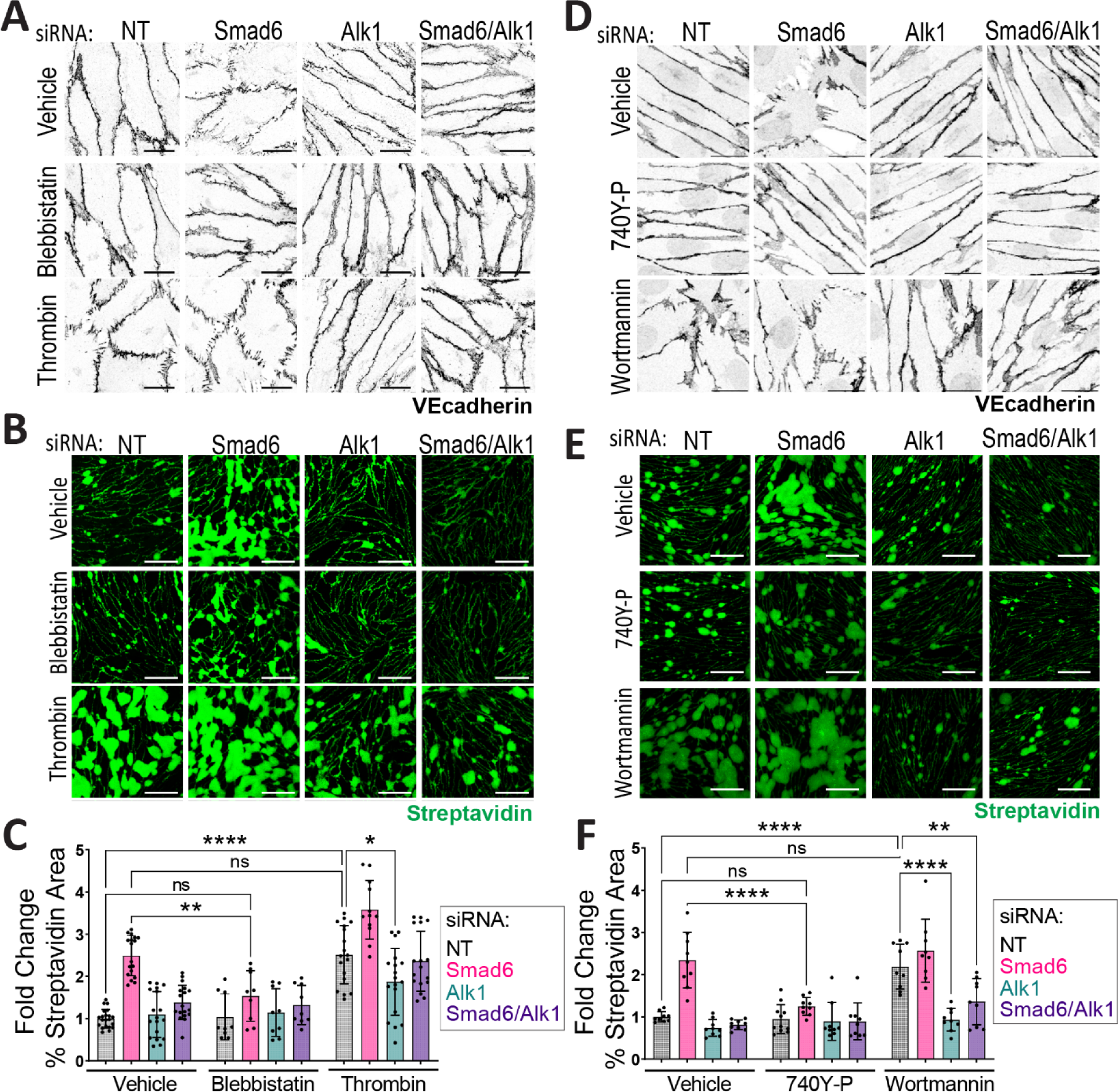
SMAD6 modulates endothelial cell hypercontractility and PI3K signaling via ALK1. HUVEC treated with non-targeting (NT), Smad6, and/or Alk1 siRNA were cultured on biotinylated fibronectin to confluence and treated as indicated for 15 min (thrombin, blebbistatin) or 21hr (740-P, wortmannin) at 37°C. **(A)** VEcadherin stain. Scale bar, 20µm. **(B)** Streptavidin-488 label. Scale bar, 100µm. **(C)** Quantification of % streptavidin-488 area/FOV (field of view). Data are mean ± SD (each data point one FOV), n = 3 experimental replicates/condition. **(D)** VEcadherin stain. Scale bar, 20µm. **(E)** Streptavidin-488 label. Scale bar, 100µm. **(F)** Quantification of % streptavidin-488 area/FOV. Data are mean ± SD (each data point one field of view), n = 3 experimental replicates per condition. *, P<0.05; **, P<0.01; ****, P<0.0001. Statistics, one-way ANOVA with Tukey’s test to correct for multiple comparisons.

ALK1 activation via BMP9 inhibits PI3K (PI3 kinase) signaling in endothelial cells (Ola et al., 2016; Alsina-Sanchís et al., 2018; Ola et al., 2018), consistent with our finding that ALK1 regulates endothelial cell contractility. We thus hypothesized that SMAD6 acts to negatively modulate ALK1 activity and maintain appropriate levels of PI3K signaling. Endothelial cell exposure to the PI3K agonist 740Y-P rescued both junction morphology and biotin matrix labeling in Smad6 depleted cells **(Fig. 8D-F)**. Conversely, inhibition of PI3K signaling via wortmannin treatment induced destabilized junction morphology and increased biotin matrix labeling in control endothelial cells to similar levels as Smad6 depletion, and Alk1 depletion blunted endothelial cell responses to wortmannin **(Fig. 8D-F)**. Thus, our results show that endothelial cell SMAD6 maintains a balance of PI3K signaling through negative modulation of ALK1 to regulate endothelial cell contractility and vessel integrity.

## DISCUSSION

Our findings reveal that SMAD6, a negative regulator of BMP signaling, is required developmentally in endothelial cells for proper blood vessel integrity. Loss of endothelial *Smad6* leads to hemorrhage and dilation of veins and capillaries of the embryonic liver, and we identify for the first time ALK1 signaling as an important negative target of *Smad6* function *in vivo*. Mechanistically, Smad6 modulates Alk1 activity to balance endothelial cell contractility that is activated by Alk1, and PI3K signaling that is normally repressed by Alk1. Thus, SMAD6 functions to maintain a balance of ALK1 signaling that in turn sets PI3K signaling levels and contractility in endothelial cells and developing blood vessels; this balance is required for vessel integrity and function **(Figure 9)** and identifies vascular ALK1 signaling as a finely tuned pathway regulated by SMAD6.

**Figure 9.**
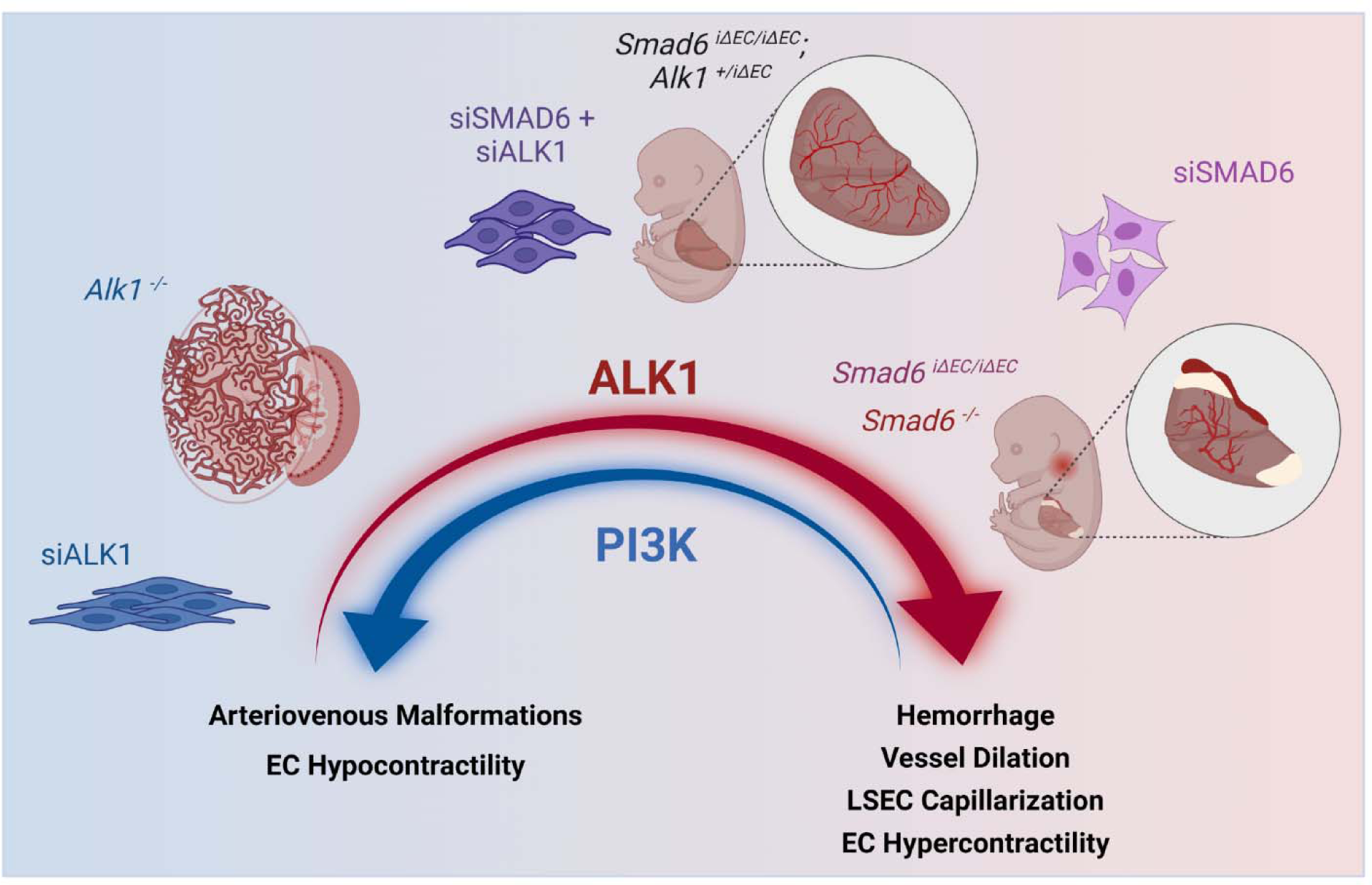
Model of SMAD6-dependent balanced ALK1 signaling to regulate endothelial cell properties and vascular development. Gain-or loss-of-function of *Alk1* leads to vascular dysfunction but with distinct phenotypes, and reduced *Alk1* gene dosage or depletion partially rescues the gain-of-function phenotypes induced by loss of the negative regulator *Smad6*. These findings identify ALK1 as a “Goldilocks” pathway in vascular development.

Despite robust SMAD6 expression in larger arteries during development (Galvin et al., 2000; Wylie et al., 2018), we found that vascular phenotypes resulting from global and endothelial-selective *Smad6* deletion were predominant in vessels characterized as veins or capillaries during embryogenesis. Our focused analysis of the embryonic liver, which receives oxygenated blood from the placenta via the umbilical vein (Swartley et al., 2016; Ober and Lemaigre, 2018), revealed significant vessel dilation and hemorrhage that likely contributed to embryonic lethality. Transcriptional profiling showed that most liver endothelial cell clusters had significant Smad6-expressing cells at both embryonic and post-natal stages (Gómez-Salinero et al., 2022). Additionally, *Smad6* was identified among the top-enriched genes in liver endothelial cells from the Tabula Muris database (Gómez-Salinero et al., 2022), consistent with a requirement for *Smad6* in embryonic endothelial cells to regulate vessel integrity. Thus, a primary developmental function of Smad6 *in vivo* is to regulate vessel function in some veins/capillaries.

Genetic loss of *Smad6* in embryonic endothelial cells resulted in a mosaic pattern of vascular loss in the in the liver parenchyma, as areas completely devoid of capillaries were juxtaposed with areas containing capillaries. As predicted, areas of mutant livers lacking capillaries appeared pale and harbored cells with more pyknotic nuclei and elevated apoptosis levels, while vascularized areas had significantly dilated capillaries that appeared intact and patent, although less mature along the LSEC differentiation lineage. Areas of hemorrhage were found within and near the avascular regions, suggesting that hemorrhage preceded capillary loss. Although the severity of the liver phenotype inversely correlated with overall Smad6 RNA levels in enriched endothelial cell populations from the same livers, the mosaic pattern of parenchymal vessel loss was also documented in embryos globally deleted for *Smad6*, indicating that the mosaicism likely results from some aspect of Smad6 function that is unique to liver vascular development.

The embryonic liver vascular phenotypes - hemorrhage, mosaic capillary loss and vessel dilation - also exhibited a striking distribution to the periphery of the liver lobes, with the central areas being relatively unaffected. This pattern may result from the developmental program of the embryonic liver, as liver sinusoids are more proliferative at the periphery (Swartley et al., 2016), and liver lobes are perfused in a peripheral to central wave developmentally (Lorenz et al., 2018). Thus, the requirement for *Smad6* function correlates with endothelial proliferation and vascular perfusion in the embryonic liver, consistent with the role of SMAD6 in flow-mediated responses of endothelial cells (Ruter et al., 2021). The concept that blood flow and flow patterns influence the spatial distribution and mosaicism of the endothelial Smad6 loss phenotype is compelling. Based on computational modeling (Bernabeu et al., 2014; Mut et al., 2009; Balogh and Bagchi, 2019; Rani et al., 2006; Hewlin and Tindall, 2023) shear stress is predicted to be highest at the point of entry of afferent vessels into an organ, and in the embryonic liver this is the umbilical/portal circulation that delivers oxygenated blood from the placenta. Thus, this venous circulation can be considered “arterialized” (Spurway et al., 2012; Swartley et al., 2016) and may exhibit a different combination of forces and molecular requirements than other vessels whose specification and function match, thus sensitizing them to *Smad6* loss. The mosaic pattern of vessel loss with *Smad6* deletion suggests that vessel rupture and hemorrhage in the umbilical/portal circulation leads to localized loss of oxygenated blood in downstream capillaries and resulting tissue deterioration. It is unclear whether the relatively mild changes to the remaining mutant capillaries are downstream of potential systemic changes or endothelial cell-autonomous results of *Smad6* loss in capillaries.

The liver is the site of synthesis of BMP9, which is a primary secreted ligand for the ALK1 arm of the BMP signaling pathway (Larrivée et al., 2012; David et al., 2007a; Miller et al., 2000), and BMP9 expression increases with developmental age through postnatal day15 (Bidart et al., 2012). This relationship led us to hypothesize that *Smad6* genetically interacts with *Alk1*, and we confirmed that BMP9 exposure mimics Smad6 depletion-induced activation of endothelial cell junctions, and both responses are Alk1-dependent, indicating that Smad6 regulates BMP9-mediated Alk1 signaling. Our results define *Alk1* as a primary target of *Smad6* in endothelial cells *in vivo*, as reduced *Alk1* dosage in endothelial cells significantly rescued the severity of the liver vascular phenotype seen with endothelial-selective *Smad6* loss, while genetic epistasis with ACVR1/Alk2 or BMPR1A/Alk3 did not show a similar relationship (data not shown), thus linking vascular *Smad6* function to the Alk1 arm of the BMP pathway. Moreover, the effects of Smad6 depletion also require Alk1 function in primary endothelial cells, as concomitant depletion of Alk1 and Smad6 rescued the loss of endothelial cell flow alignment, junction stabilization and monolayer integrity seen with Smad6 depletion alone.

Both *Smad6* and *Alk1* appear required for vessel integrity and vascular homeostasis, since embryonic endothelial-selective deletion of each gene is accompanied by vascular hemorrhage and lethality (Wylie et al., 2018; Park et al., 2008b). This apparent paradox of a negative regulator having a similar phenotype to its target suggests a more complex relationship, and several lines of evidence suggest that the balance of Alk1 activity in endothelial cells may be crucial to proper blood vessel integrity. Although both Smad6 and Alk1 are required for endothelial cell flow responses, the outputs of depletion in endothelial cells are different, with Smad6 depletion leading to misalignment while Alk1 depletion results in hyper-alignment *in vitro* ((Ruter et al., 2021); this study). Loss of Alk1 is associated with arterio-venous malformations (AVMs) *in vivo* (Park et al., 2021; Urness et al., 2000; Tual-Chalot et al., 2014; Park et al., 2009), while Smad6 loss is not associated with AVMs but rather with perturbed barrier function and hemorrhage. These findings suggest distinct functions for SMAD6 and ALK1 in endothelial flow responses. Here we show that endothelial cell contractility is negatively regulated by SMAD6 but positively regulated by ALK1, and PI3K signaling is likely positively regulated by SMAD6, while others have shown that ALK1 signaling negatively regulates PI3K signaling (Ola et al., 2016; Jin et al., 2017). These findings show that balanced Alk1 signaling normally leads to the proper level of PI3K signaling and endothelial cell contractility, and that this balance is regulated by *Smad6* for proper endothelial cell flow responses and vessel integrity important in liver vascular development.

More broadly, our findings suggest that the role of Alk1 signaling in transducing vascular flow responses is complex and nuanced in endothelial cells, with loss-of-function leading to inappropriate flow responses (Poduri et al., 2017; Peacock et al., 2020) while the proposed gain-of-function in Alk1 signaling also affects flow responses in different ways (Ruter et al., 2021). A major feature of the gain-of-function phenotype revealed by SMAD6 loss is hyper-contractility associated with reduced PI3K signaling, which is consistent with the effects of SMAD6 on endothelial junctions (Wylie et al., 2018). Taken together, these findings suggest that endothelial Alk1 signaling is an example of a “Goldilocks” pathway that requires a certain signaling amplitude to function properly, similar to the regulation described for cytokine signaling and neural circuits (Graham et al., 2022; Humphries, 2016; Petersen and Berg, 2016). Moreover, Smad6 is a prominent transcriptional target that is upregulated downstream of BMP9/Alk1 signaling, consistent with the idea that negative modulation via SMAD6 is important to counteract positive inputs and promote vessel integrity. Current therapies to mitigate symptoms of HHT2, a disease resulting from genetic loss of the Alk1 arm of BMP signaling, include PI3K blockade (Robert et al., 2020); our work suggests that modulating the Alk1 arm of BMP signaling may restore the required balance in the pathway.

## MATERIALS AND METHODS

### Mice

All animal experiments were approved by the University of North Carolina at Chapel Hill (UNC-CH) Institutional Animal Care and Use Committee. All mice were on a C57BL/6J genetic background, and both male and female embryos were included. *Smad6^+/-^* mice (Galvin et al., 2000; Wylie et al., 2018) were backcrossed to the C57BL/6J background for N≥10 generations. *Cdh5Cre^ERT2^* (Tg(Cdh5-cre/ERT2)1Rha) mice (Sörensen et al., 2009) were obtained from Cancer Research UK. *UBC-Cre^ERT2^* mice (B6.Cg-Ndor1Tg(UBC-cre/ERT2)1Ejb/2J) were previously described (Ruzankina et al., 2007). *Smad6^fl/+^* mice were generated by the UNC-CH Animal Models Core via introduction of LoxP sites around exon 4 of the *Smad6* gene. To induce genetic deletion, tamoxifen (Sigma, T5648) in sunflower oil was administered to timed-pregnant dams on day E10.5 via oral gavage at 0.12mg/g body weight (Park et al., 2008a). *Cre^ERT2^* negative littermates were used as controls. Embryos were collected at indicated timepoints into PBS on ice, euthanized according to IACUC approved methods, and fixed in 4% paraformaldehyde (PFA) at 4°C for 24-72 hr.

For DNA analysis, embryonic tail snips or lung tissue was incubated in 0.2mg/mL Proteinase K in DirectPCR Lysis Reagent (Viagen Biotech 101-T) at 55°C for 4 hr, followed by enzyme inactivation at 85°C for 45 min. Forward and reverse primers (Smad6 excised F+R) were designed to anneal upstream of the 5’ loxP site and downstream of the 3’ loxP site (**Supp. Fig 1B**). See **Table 1** for primer details.

**Table 1.**
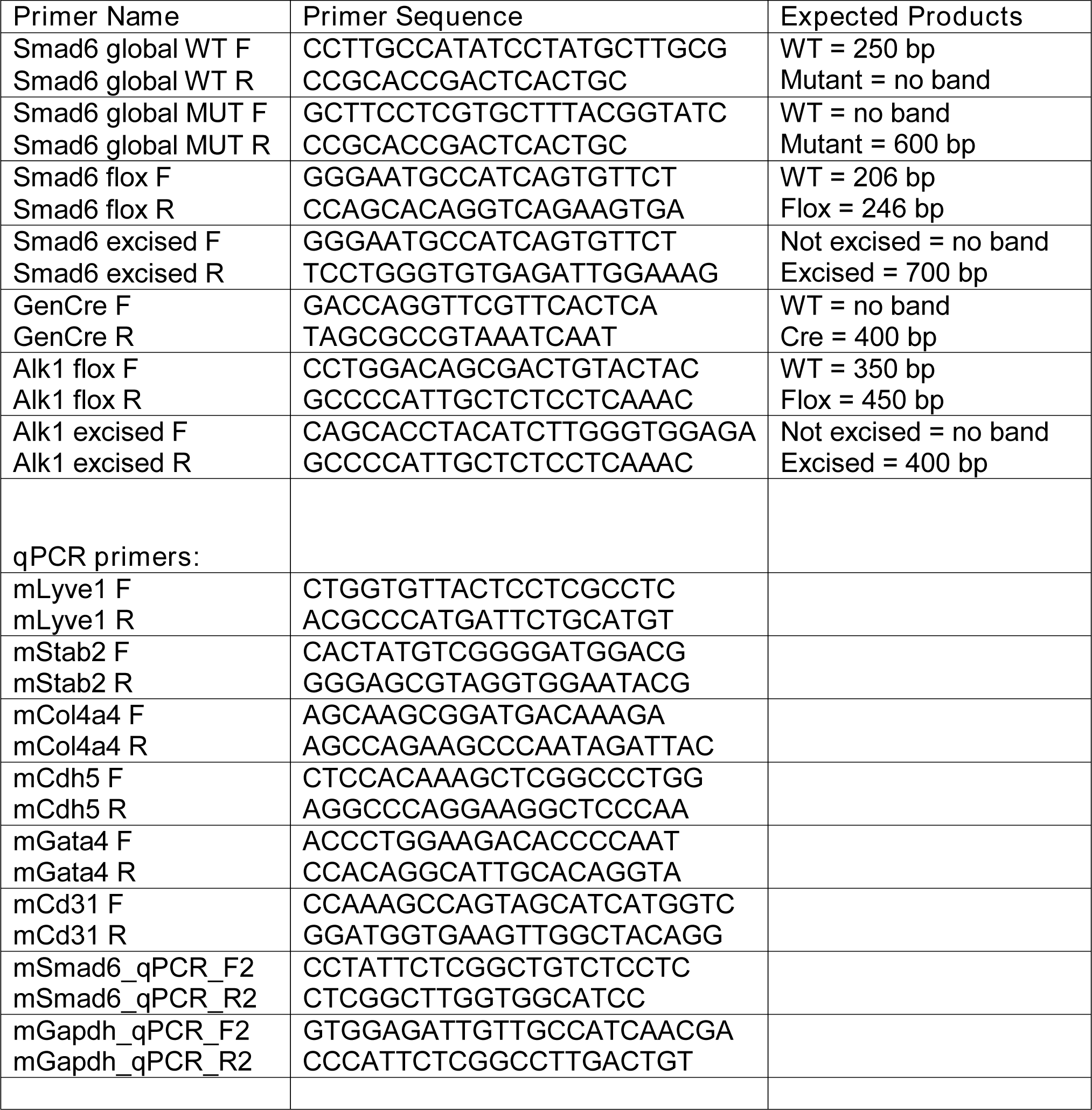
PCR Primers

### MACS Enrichment and qRT-PCR

This protocol was adapted from (Sokol et al., 2021). Livers isolated from E16.5 embryos were minced and digested in 6mL of digestion buffer (250U/mL Collagenase type II (Worthington LS004204), 0.6U/mL Dispase (Worthington LS02104), 33U/mL DNAse I (USB/Affymetrix 14340), in EBM basal media (Lonza NC1447083)) at 37°C for 30 min with gentle vortexing. Cell suspensions were broken up by passage thru a 18G1-1/2 needle linked to a 20mL syringe, neutralized in 8mL EGM-2 (Lonza NC9525043) + 20% NBCS (Gibco 16010-159), then centrifuged at 300x g for 7min. Supernatant was removed and cells resuspended in 1mL RBC Lysis Buffer (Miltenyi 130-094-183) for 2 min. Following a spin at 300x g for 5 min, cells were resuspended in 5mL cold buffer (DPBS + 2% NBCS + 2mM EDTA), spun again at 300x g spin for 7min, then resuspended in ice-cold MACS buffer (DPBS + 0.5% BSA + 2mM EDTA) and incubated with Fc Block (Biolegend 156603) for 5 min at 4°C. Cells were mixed with PECAM1 (CD31) antibody-labeled beads (Miltenyi 130-097-418) at 4°C for 15 min, washed in cold MACS buffer, then resuspended in MACS buffer and passed through MS columns (Miltenyi 130-042-201) on a MACS MultiStand Separator. Flow-through was collected, then columns removed from the stand and washed to collect PECAM1+ endothelial fraction. Cells were pelleted and resuspended in Trizol at −80°C. RNA isolation was using the Direct-Zol RNA MiniPrep kit (Zymo, R2052), followed by iScript cDNA Synthesis (BioRad 1708891) according to manufacturer instructions. RT-qPCR was run on a QuantStudio 6 Flex Real-Time PCR system (Applied Biosystems) with iTaq Universal SYBR Green Supermix (BioRad 1725121) and primers outlined in **Table 1**. Results were analyzed using delta-delta-CT methods, and CT values were normalized to GAPDH or B-actin and relative to the WT average.

### Semi-Quantitative Phenotype Score

Intact embryos were examined and imaged. For liver phenotype scores, livers were dissected either pre-or post-fixation and imaged. A severity guideline key (**Supp. Fig 2**) was created that ranked presentations of each phenotype (jugular hemorrhage, abdominal hemorrhage, liver hemorrhage/paleness). Embryo genotypes were blinded and images scored by a single researcher to maintain internal consistency.

### LacZ Staining

LacZ staining was performed as described (Nagy et al., 2007) with modifications as follows:

#### Wholemount

Briefly, E16.5 embryos in PBS on ice were dissected and the heart, lungs, liver, and intestines removed. All tissues were incubated in freshly made 0.2% glutaraldehyde (Electron Microscopy Sciences cat no. 16120) + 5mM EGTA + 2mM MgCl_2_ in PBS on ice for 30 min, washed in wash buffer (2mM MgCl_2_ + 0.02% IGEPAL + 0.01% sodium deoxycholate in 0.1M Sodium Phosphate Buffer pH 7.3) for 3x 15 min at RT, then incubated in freshly made stain solution (5mM potassium ferricyanide + 5mM potassium ferrocyanide + 1mg/mL X-gal (Promega cat no. V3941) in wash buffer) for 8 hr at 37°C with gentle rocking. Tissues were washed in PBS, then incubated in 4% PFA in PBS for 1hr at RT before imaging with a stereomicroscope or embedding in paraffin.

#### Frozen Sections

E16.5 embryos were euthanized, rinsed in PBS with Mg^2+^ and separated above the liver. Embryos were fixed in cold 0.25% glutaraldehyde (Electron Microscopy Sciences cat no. 16120) in PBS, washed 3×5 min in PBS, and sunk in 30% sucrose in PBS at 4°C for 12hr. The embryo pieces were embedded in OCT, frozen, sectioned at 10µm, and stored at −80°C. Before staining, sections were warmed at room temp for 20min, washed in PBS 5min, fixed in 0.25% glutaraldehyde in PBS 5min at room temp, washed 3×5min in PBS. Slides were incubated in freshly made stain solution (5mM potassium ferricyanide + 5mM potassium ferrocyanide + 1mg/mL X-gal (Promega cat no. V3941) in wash buffer) overnight at 37°C. Sections were post-fixed in 4% PFA for 1hr at RT, washed in PBS 3×5min, counterstained with Nuclear Fast Red (Sigma, N3030) 4 min, and mounted in 80% glycerol.

### Histology and Immunofluorescence

#### H&E (hematoxylin and eosin)

Fixed tissues were embedded in paraffin, sectioned at 10µm thickness, deparaffinized in 2x xylene washes (Fisher Chemical cat no. X3S-4) for 10 min, and rehydrated in gradients of ethanol (100%, 95%, 70%) to pure dH_2_O. H&E stain was as described in (Cardiff et al., 2014). Briefly, sections were incubated in acidified Harris hematoxylin (Thermo Fisher 6765003) for 8 min, rinsed in dH_2_O, incubated in 1% acid alcohol 30 sec, rinsed, put in Bluing Reagent (Fisher Chemical cat no. 220-106) 30 sec, rinsed, stained with Eosin Y Alcohol (0.25% Eosin Y (Fisher Chemical SE23-500D) in 80% ethanol and 0.5% glacial acetic acid) 4 min, then dehydrated in 100% ethanol 2x 30 sec, incubated in xylene, and mounted with OmniMount (National Diagnostics cat no. HS-110).

#### Immunofluorescence

For formalin-fixed paraffin-embedded (FFPE) sections, samples were deparaffinized and rehydrated as above. Frozen sections (described above) were set out at RT for 20 min followed by rehydration in PBS for 20 min. Antigen retrieval was performed in citrate buffer (pH 6.0) (Vector Labs cat no. H-3300) in a steamer for 40 min for FFPE sections, or 5 min for frozen sections. Slides were cooled at RT 20 min, washed in PBS, then permeabilized in 0.1% Triton-X in PBS (PBSTx) 15 min at RT. Sections were blocked in 5% normal donkey serum (Sigma cat no. D9663) in 0.1% PBSTx for 1 hr at RT. Unconjugated primary antibodies were diluted in block according (see **Table 2**) and incubated overnight at 4°C. Sections were washed in PBS, re-blocked 20 min at RT, then incubated in secondary antibodies, DAPI, and fluorescently-conjugated primary antibodies (see **Table 2**). Slides were rinsed in PBS, then mounted in Prolong Diamond Antifade mounting media (Life Technologies cat no. P36961).

**Table 2.** Antibodies

| **Antibody** | **Species** | **Company** | **Catalog No.** | **Dilution** |
| --- | --- | --- | --- | --- |
| LYVE-1 | Goat | R&D Biosystems | AF2125 | 1:100 |
| PECAM (CD31) | Goat | R&D Biosystems | AF3628 | 1:50 |
| VEcadherin | Goat | R&D Biosystems | AF1002 | 1:50 |
| VEGFR3 | Goat | R&D Biosystems | AF743 | 1:50 |
| ERG | Rabbit | Abcam | Ab92513 | 1:100 |
| Ter119 | Rat | R&D Biosystems | MAB1125 | 1:100 |
| Cleaved Caspase 3 | Rabbit | Cell Signaling Technology | 9661 | 1:400 |
| Collagen IV | Rabbit | GeneTex | GTX19808 | 1:200 |
| αSMA-Cy3 | n/a | Sigma | C6198 | 1:500 |
| DAPI |  | Sigma | 10236276001 | 1:1,000 |
| Donkey-α-Goat 647 | | Life Technologies | A-21447 | 1:250 |
| Donkey-α-Goat 488 | | Life Technologies | A-11055 | 1:250 |
| Donkey-α-Goat 594 | | Life Technologies | A-11058 | 1:250 |
| Donkey-α-Rat 488 | | Life Technologies | A-21208 | 1:250 |
| Donkey-α-Rabbit 488 | | Life Technologies | A-21206 | 1:250 |
| Donkey-α-Rabbit 594 | | Life Technologies | A-21207 | 1:250 |
| Donkey-α-Rabbit 647 | | Life Technologies | A-31573 | 1:250 |

\*Antibodies validated using secondary-only controls, and specificity was confirmed by staining tissues/structures known to be positive.

Following fixation in 4% PFA, HUVEC were washed with PBS, permeabilized in 0.1% Triton X-100 (Sigma T8787) at RT for 10 min and blocked at RT for 1 hr in blocking solution (5% NBCS, 2x antibiotic-antimycotic (Gibco), 0.1% sodium azide (Sigma s2002-100G). Cells were incubated in primary antibody **(Table 2)** diluted in blocking solution at 4°C overnight and washed in PBS 3x 15 min. Secondary antibodies **(Table 2)** with DAPI were diluted in blocking solution and added for 1 hr at RT, then washed in 3x 10 min PBS. Slides were mounted with coverslips and Prolong Diamond Antifade mounting medium (Life Technologies P36961).

### Single Cell RNA Sequencing Analysis

#### Mouse Organogenesis Cell Atlas (Cao et al., 2019)

Analysis was performed using the R package Seurat. The MOCA dataset contains over 2 million cells and the gene count matrix is over 20GB, so a randomly down-sampled dataset containing 10,000 cells was downloaded for QC check and EC annotation. Dimension reduction results from t-distributed stochastic neighbor embedding (t-SNE) and QC showed that cell clusters did not correlate with total detectable molecules/cell (nCount_RNA) or the number of detectable genes/cell (nFeature_RNA), suggesting that data from MOCA were analyzed properly. Next, cell type annotations in the dataset and expression patterns of pan-endothelial markers Cdh5 and Pecam1 were plotted. Only clusters annotated as EC and endocardial cells show high levels of Cdh5 and Pecam1, indicating that annotation in the data is correct. After confirming that QC was properly performed and EC annotation was correct in MOCA with this subset of data (data not shown), EC were extracted from the original gene count matrix and subjected to t-SNE visualization to show EC clusters labeled by developmental stage (**Fig 2A**) and inferred embryonic tissue origin (**Fig2 A’**). The expression of genes of interest was plotted by FeaturePlot (**Fig.2 A’’**) and VlnPlot.

#### Fetal Liver (Gómez-Salinero et al., 2022)

The R object GSE174209_RObject_Timepoints.Rdata was downloaded from the Gene Expression Omnibus (https://www.ncbi.nlm.nih.gov/geo/query/acc.cgi?acc=GSE174209) and analyzed using Seurat. Fetal EC were extracted from the original gene count matrix (post-natal EC were excluded) and subjected to UMAP visualization to show EC clusters labeled by different fetal liver endothelial cell populations. UMAP and violin plots were generated to highlight SMAD6 expression in these fetal liver EC populations.

#### EC Atlas of Adult Mouse Tissues (Kalucka et al., 2020)

We reanalyzed the published single-cell (sc) RNA seq dataset from an EC Atlas of adult mouse tissues via their online portal (https://endotheliomics.shinyapps.io/ec_atlas/) (**Sup. Fig4 A-D’**). In the portal we selected liver, lung, and brain as tissue sets of interest, and searched “Smad6” to auto-generate t-SNE plots of Smad6 expression.

#### Adult Mouse Brain & Lung Vascular Cells (Vanlandewijck et al., 2018; He et al., 2018)

We reanalyzed the published single-cell (sc) RNA seq dataset from an adult mouse brain and lung vascular/perivascular scRNA seq dataset via their online portal (http://betsholtzlab.org/VascularSingleCells/database.html) (**Sup. Fig4 E-F**). In the portal we searched “Smad6” and reported the average counts of vascular endothelial populations from the auto-generated graph.

### Cell Culture and Treatments

Human umbilical vein endothelial cells (HUVEC, Lonza C2519A) were cultured at 37°C and 5% CO_2_ in EGM^TM^ Endothelial Cell Growth Medium with BulletKit^TM^ (Lonza CC-3124) and 1x antibiotic-antimycotic (Gibco) and used prior to passage 7. For contractility assays, HUVEC were treated with 0.5U/mL thrombin (Sigma T7201-500UN) at 37°C for 15 min. For contractility inhibition assays, HUVEC were treated with 10µM blebbistatin (Sigma B0560-1MG) at 37°C for 15 min. For PI3K activation assays, HUVEC were treated with 20µM 740Y-P (MedChemExpress HY-P0175) at 37°C for 22 hr. For PI3K inhibition assays, HUVEC were treated with 100nM wortmannin (SelleckChem, S2758) at 37°C for 22 hr. For BMP9 ligand assays, HUVEC were serum starved in Endothelial Base Media (Lonza CC-3162) with 0.1% FBS for 24 hr followed by treatment with 10ng/mL BMP9 (R&D Systems 3209-BP-010) at 37°C for 1 hr. Immediately following all drug treatments, HUVEC were fixed in warm 4% PFA at RT for 4 min.

### siRNA Depletion

HUVEC were transfected with non-targeting siRNA (NT) (Silencer Select Negative Control #2 siRNA, Life Technologies 4390847), SMAD6 (SMARTpool ON-TARGETplus Human SMAD6 siRNA, Dharmacon L-015362-00-0005) or (SMAD6 siRNA pool, Santa Cruz Biotechnology, sc-38380), and/or ALK1 (SMARTpool ON-TARGETplus Human ACVRL1 siRNA, Dharmacon L-005302-02-0005) using Lipofectamine 3000 (ThermoFisher L3000015) according to manufacturer directions. HUVEC were transfected at 70% confluence for 24 hr at 37°C, then incubated with fresh EGM-2 for 24 hr. Cells were seeded onto glass chamber slides coated with 5µg/mL fibronectin (Sigma, F2006-2MG) for experiments.

### Endothelial Cell Flow Experiments

Flow experiments were performed using an Ibidi pump system as described (Ruter et al., 2021) with adjustments as follows: HUVEC were seeded onto fibronectin-coated Ibidi slides (µ-Slide I Luer I 0.6mm (catalog #80186), or µ-Slide Y-shaped (catalog #80126)) in flow medium (EBM-2 with 2% FBS, 1x antibiotic-antimycotic, and 1% Nyastatin) at a density of 2×10^5^ cells/mm^2^. The next day HUVEC were exposed to 7.5 dyn/cm^2^ laminar shear stress for 72 hr.

### Biotin Matrix-Labeling

Labeling of biotinylated matrix was modified from (Dubrovskyi et al., 2013). Briefly, 0.1mg/mL fibronectin was incubated with 0.5mM EZ-Link Sulfo-NHS-LC-Biotin (ThermoFisher A39257) for 30 min at RT. Biotinylated fibronectin (0.5µg/mL) was coated onto glass chamber slides for 30 min at RT, then HUVEC were seeded at a density of 7.5×10^4^ cells/mm^2^. Following drug treatments or flow experiments, confluent HUVEC were treated with 25µg/mL Streptavidin-488 (Invitrogen S11223) for 3 min at RT then immediately fixed in warm 4% PFA as described above. For quantification, at least three 40x confocal z-stack images/condition/experiment were taken. The streptavidin channel was thresholded in ImageJ and the % labeled area measured, then normalized to the siNT control average for each respective experiment.

### Real Time Cell Analysis (RTCA)

An xCELLigence Real-Time Cell Analyzer (RTCA, Acea Biosciences/Roche Applied Science) was used to assess barrier function of HUVEC monolayers. HUVEC were seeded at a density of 60,000cells/well of the E-plate (E-plate 16, Roche Applied Science), then electrical impedance readings acquired every 2 min for 24 hr. Results are reported at 24 hr as the percent change in cell index calculated using the following formula: (Cell Index_siRNA_ – Cell Index_NT_)/ABS(Cell Index_NT_).

### Imaging and Analysis

Whole embryo and intact liver images were acquired using a Leica MZ 16 F stereomicroscope and an Olympus DP71 camera. H&E stains were scanned at 20x on an Olympus SLIDEVIEW VS200. Images of fluorescently stained tissue sections were acquired using an Olympus FV300 confocal microscope with Fluoview software or on an Olympus SLIDEVIEW VS200 with OlyVIA software. Images were processed in ImageJ or QuPath software and shown in figures as compressed Z stacks.

#### Vascularized liver area

Scans of whole liver sections were traced along the outer edge of the DAPI channel to measure total liver area. Traces were then made on the LYVE1+ channel around the vascularized zones. <u>Apoptosis:</u> Scans of whole liver sections were imported into QuPath, and the Cleaved Caspase 3 channel was thresholded and measured for positive staining area in µm^2^. <u>Collagen IV:</u> 40x compressed Z-stack images were thresholded and measured for positive staining area relative to image area. At least 4 images/embryo were taken from similar regions in each liver. <u>Capillary dilation:</u> representative 40x compressed Z stack images were obtained in capillary beds along the edges of the livers. Lines were drawn in ImageJ perpendicular to each capillary to measure diameter at the widest point between branchpoints. <u>Cell elongation under flow:</u> HUVEC stained with PECAM1 were measured and the longest axis of cell divided by shortest axis. At least four 40x compressed Z-stack images were measured per condition.

### Statistical analysis

X^2^ analyses were run for categorical data (expected vs. observed genotype and for semi-qualitative phenotype comparison graphs). GraphPad Prism 9.4.1 software was used to perform all other statistical comparisons, and all comparisons were two-tailed. In experiments with two groups, Student’s two-tailed unpaired *t-*test was used to determine statistical significance. For % change in RTCA experiments a one sample t-test against a hypothetical mean of “0” was used. One-way ANOVA with Tukey’s test to correct for multiple comparisons was used to compare differences between more than two groups. For thrombin, blebbistatin, 740Y-P, and wortmannin experiments, two-way ANOVA with Tukey’s multiple comparisons test was used to determine statistical significance.

## ACKNOWLEDGEMENTS

We thank the UNC Animal Models Core for assistance in generating the *Smad6* floxed mouse line. We thank Wendy Salmon (Hooker Imaging Core, UNC) for microscopy support. The UNC Hooker Imaging Core Facility is supported in part by P30 CA016086 Cancer Center Core Support Grant to the UNC Lineberger Comprehensive Cancer Center. We thank Caroline Crater for mouse room support, and Bautch Lab members for critical discussion and feedback. We thank the Center for Gastrointestinal Biology and Disease core at UNC for histology support. We acknowledge use of the publicly available scRNAseq databases from the Rafii lab (embryonic and post-natal liver), the Shendure lab (Mouse Organogenesis Cell Atlas), the Carmeliet lab (EC Mouse Atlas), and Betsholtz lab (Lung and Brain EC Atlas).

## COMPETING INTERESTS

None.

## FUNDING

This work was supported by grants from the National Institutes of Health-NHLBI (R35 HL139950 to VLB), and the National Science Foundation Graduate Research Fellowship Program (DGE-1650116 to MRK).

## DATA AVAILABILITY

RNA seq analyses presented are from previously published publicly available sources (Cao et al., 2019; Gómez-Salinero et al., 2022; Kalucka et al., 2020; Vanlandewijck et al., 2018; He et al., 2018). Other data is presented in the manuscript or available upon reasonable request.

## SUPPLEMENTAL FIGURE LEGENDS

**Supplementary Figure 1. Characterization of *Smad6^-/-^* and *Smad6^i^*^Δ^*^EC/i^*^Δ^*^EC^* Mice. (A)** C57BL/6J *Smad6^+/-^* x *Smad6^+/-^* intercrosses genotyped at indicated embryonic stages or P0 (non-viable embryos excluded). Green shading, significantly reduced numbers of *Smad6^-/-^* P0 pups; purple shading, embryonic stage (E16.5) used for phenotype analyses. ****, P<0.0001. Statistics, Χ^2^ analysis. **(B)** Schematic for generation of *Smad6* floxed allele. LoxP sites were engineered around Exon 4 (encoding a functional MH2 domain) of the *Smad6* gene. Primers surrounding the 5’ LoxP site (F1 and R1) were used to confirm presence of floxed allele, while primers F1 and R2 were used to determine excision. **(C)** Breeding and excision/collection scheme for *Smad6^i^*^Δ^*^EC/i^*^Δ^*^EC^* embryos/pups. **(D)** C57BL/6J *Smad6^fl/fl^;Cdh5-Cre^ERT2/+^* x *Smad6^fl/fl^* intercrosses genotyped at P0 or E16.5 (non-viable pups excluded). *, P<0.01. Statistics, Χ^2^ analysis.

**Supplementary Figure 2. Semi-Quantitative Phenotype Scoring Key for E16.5 Embryos/Livers. (A)** E16.5 embryo images illustrate scoring criteria for phenotypic severity for indicated hemorrhage areas. Arrowheads, location of hemorrhage. **(B)** Isolated E16.5 livers illustrate scoring criteria for phenotypic severity. Asterisks, pale regions; arrows, hemorrhage/dilated vessels.

**Supplementary Figure 3. *Smad6^UBCCreER^* Embryo Characterization & *Smad6^i^*^Δ^*^EC/i^*^Δ^*^EC^* Excision Efficiency. (A)** Breeding scheme and excision/collection schedule for *Smad6^i^*^Δ^*^/i^*^Δ^ mice. **(B)** Representative images of *Smad6^i^*^Δ^*^/i^*^Δ^ E16.5 embryos and control. Scale bar, 2.5mm. **(C-E)** Semi-quantitative whole embryo analysis of indicated genotypes (see **Supp. Fig 2A** for criteria). Categories: total hemorrhage (C); abdominal hemorrhage **(D)**; jugular hemorrhage **(E)**. WT, n=18 embryos; *Smad6^i^*^Δ^*^/i^*^Δ^, n=10 embryos. **(F)** Representative images of E16.5 *Smad6^i^*^Δ^*^/i^*^Δ^ livers and control. Asterisks, pale regions. Arrows, vascular dilation/hemorrhage. Scale bar, 1mm. **(G)** Semi-quantitative phenotype analysis of isolated livers (see **Supp. Fig. 2B** for criteria), WT, n=8 livers; *Smad6^i^*^Δ^*^/i^*^Δ^, n=5 livers. ***, P<0.001; ns, not significant. Statistics, Χ^2^ analysis. **(H)** Breeding scheme and schedule for *Smad6^i^*^Δ^*^EC/i^*^Δ^*^EC^* mice in (I-K). **(I)** PCR analysis of E16.5 embryo lung DNA of indicated genotypes. (J) qRT-PCR for Smad6 RNA in PECAM1-enriched or PECAM1 negative cells from E16.5 livers of indicated genotypes. CT values normalized to Gapdh. mRNA fold change relative to WT average. WT, n=6 livers; *Smad6^i^*^Δ^*^EC/i^*^Δ^*^EC^*, n=8 livers. **, P<0.01; ns, not significant. Data, individual data point/embryo ±SD. Statistics, one-way ANOVA with Tukey’s test to correct for multiple comparisons. **(K)** Liver phenotype score relative to fold change Smad6 mRNA, with linear regression. R^2^ = 0.9015; P=0.0003. *Smad6^i^*^Δ^*^EC/i^*^Δ^*^EC^*, n=6 livers.

**Supplementary Figure 4. Smad6 is Expressed in Mouse Brain, Lung, & Liver Endothelial Cells. (A-D)** scRNAseq data from EC (endothelial cell) Atlas of 11 tissues from adult mice (https://endotheliomics.shinyapps.io/ec_atlas/)(Kalucka et al., 2020). **(A)** All tissues clustered by organ source, **(B-D)** individual organ EC clusters. (A’) UMAP of Smad6 expression in all tissues. (**B’-D’**) Individual organ EC clustered by endothelial subsets. **(E-F)** scRNA-seq dataset from adult mouse brain and lung endothelial cells (http://betsholtzlab.org/VascularSingleCells/database.html) (Vanlandewijck et al., 2018; He et al., 2018) **(E)** Smad6 expression by average counts in Lung EC. [Lung data]: EC - endothelial cells; capil, capillary; a, arterial; c, continuum; L, lymphatic; 1,2,3 - subtypes. (F) Smad6 expression by average counts in Brain EC. [Brain data]: EC - endothelial cells; v, venous; capil, capillary; a, arterial; aa, arteriolar; 1,2,3 - subtypes.

**Supplementary Figure 5. Embryonic lethality of endothelial *Alk1* homozygous deletion.(A)** Representative images of embryos from *Alk1^fl/+^;Cdh5Cre^ERT2/+^* x *Alk1^fl/fl^* cross with tamoxifen gavage at E10.5 and collection at E16.5. Data from 3 litters. 5 non-viable embryos were genotyped as *Alk1^fl/fl^;Cdh5Cre^ERT2/+^*. **(B)** Representative images of embryos from an *Alk1^fl/+^;Smad6^fl/fl^;Cdh5Cre^ERT2/+^* x *Alk1^fl/fl^;Smad6^fl/fl^* cross with tamoxifen gavage at E10.5 and collection at E16.5. Data from 1 litter. 1 non-viable embryo was genotyped as *Alk1^fl/fl^;Smad6^fl/fl^;Cdh5Cre^ERT2/+^*.

**Supplementary Figure 6. RNA expression of PECAM1^neg^ E16.5 liver cells** RNA isolated from PECAM1-negative fractions (PECAM1^neg^) of E16.5 *Smad6^i^*^Δ^*^EC/i^*^Δ^*^EC^* or control littermate livers. qRT-PCR for **(A)** Lyve1, **(B)** VEcadherin, and **(C)** CD31. CT values normalized to Gapdh and mRNA expression reported as fold change relative to WT average. WT, n=6; *Smad6^i^*^Δ^*^EC/i^*^Δ^*^EC^*, n=8 livers. ns, not significant. Data are individual data points for each embryo ±SD. Statistics, one-way ANOVA corrected for multiple comparisons.

**Supplementary Figure 7. BMP9 destabilizes endothelial cell junctions.** HUVEC treated with non-targeting (NT), Smad6, and/or Alk1 siRNA were cultured on fibronectin to confluence and treated as indicated with BMP9 ligand for 24 hr at 37°C. **(A)** PECAM1 and DAPI stain. Scale bar, 20µm. Representative images of n = 3 experimental replicates.

## Notes

### Competing Interest Statement

The authors have declared no competing interest.

